# Spatiotemporal patterns of rodent hippocampal field potentials uncover spatial representations

**DOI:** 10.1101/828467

**Authors:** Liang Cao, Viktor Varga, Zhe S. Chen

**Affiliations:** The Neuroscience Institute, New York University School of Medicine, New York, NY 10016, USA; Department of Physics, East China Normal University, Shanghai, China; Institute of Experimental Medicine, Budapest, Hungary; Department of Psychiatry, Department of Neuroscience and Physiology, New York University School of Medicine, New York, NY 10016, USA

## Abstract

Spatiotemporal patterns of large-scale spiking and field potentials of the rodent hippocampus encode spatial representations during maze run, immobility and sleep. Here, we showed that multi-site hippocampal field potential amplitude at ultra-high frequency band (FPA_uhf_) provides not only a fast and reliable reconstruction of the rodent’s position in wake, but also a readout of replay content during sharp wave ripples. This FPA_uhf_ feature may serve as robust real-time decoding strategy from large-scale (up to 100,000 electrodes) recordings in closed-loop experiments. Furthermore, we developed unsupervised learning approaches to extract low-dimensional spatiotemporal FPA_uhf_ features during run and ripple periods, and to infer latent dynamical structures from lower-rank FPA_uhf_ features. We also developed a novel optical flow-based method to identify propagating spatiotemporal LFP patterns from multi-site array recordings, which can be used for decoding application. Finally, we developed a prospective decoding strategy to predict animal’s future decision in goal-directed navigation.

## Introduction

Cutting-edge large-scale high-density electrode arrays have enabled us to record spatially distributed neural activity within local or multiple brain circuits (Berényi et al., 2014; Jun et al., 2017; Chung et al., 2019), yet it remains challenging to reliably uncover representations of spatiotemporal neural activity. The hippocampus contributes to a wide range of brain functions responsible for spatial and episodic memories, learning and planning (Pfeiffer and Foster, 2013). Population spike activities from hippocampal place cell assemblies provide a readout of the rat’s spatial location (Zhang et al., 1998; Davidson et al., 2009; Kloosterman et al., 2014; Grosmark and Buzsáki, 2016; Hu et al., 2018; Ciliberti et al., 2018). However, direct use of spike information for neural decoding remains challenging due to practical issues of spike sorting and unit instability, let alone the prohibitive computational cost in the era of big data (Rey et al., 2015; Rossant et al., 2016). In contrast, hippocampal field potentials consist of collective local synaptic potentials around the recording site, serving as an alternative information source for spatial representation (Buzsáki et al., 2012; Agarwal et al., 2014). In addition to the animal studies, high-density electrocorticography (ECoG) grid has allowed us to access large-scale field potentials from human brain (Khodagholy et al., 2015; Chang et al., 2015; Zhang et al., 2015). Developing effective statistical methods for uncovering neural representations of these signals during memory tasks or sleep remains a central goal in computational neuroscience.

Hippocampal sharp-wave ripples (SPW-Rs) are important hallmarks for memory reactivations during immobility and non-rapid-eye-movement (NREM) sleep (Roumis and Frank, 2015; Buzsáki et al., 2015; Fernández-Ruiz et al., 2019). Decoding the content of hippocampal replays during ripple events can help dissect circuit mechanisms of memory consolidation, planning and decision-making (Pfeiffer and Foster, 2013; Davidson et al., 2009; Foster, 2017; Grosmark and Buzsáki, 2016; Ólafsdóttir et al., 2018). Evidence has suggested that hippocampal LFPs may encode neuronal ensemble activations during SPW-Rs (Taxidis et al., 2015), but it remains unknown how the information embedded in field potentials is related to spatial representations, and how these spatially distributed hippocampal filed potential representations relate to ensemble spike representations at different brain states.

To address these questions and challenges, here we systematically investigate spatiotemporal patterns derived from large-scale multi-site recorded rodent hippocampal field potentials, and their spatial representations during maze running, memory replay decoding, and spatial decision-making. We propose a set of independent and complementary spatiotemporal features derived from hippocampal field potentials, and develop innovative supervised or unsupervised learning methods while validating them in various applications. These robust spatiotemporal features not only provide direct and robust solutions to decoding and prediction problems, but also open new opportunities for visualizing high-dimensional neural data or finding latent structures in the absence of behavioral measure.

## Results

### Spatially distributed hippocampal field potentials encode position in maze run

Rats and mice were implanted with large-scale or high-density silicon probes (varying from 64 to 512 channels; see **Figure 1 — table supplement 1**) in the dorsal hippocampus or hippocampi while freely foraging in a circular track, linear track, T-maze or open field environment (see **Figure 1 — figure supplement 1**). We processed the extracellular raw voltage traces of hippocampal recordings and extracted independent neural signals at different frequency bands (**Figure 1a**). During run epochs, we found that both demodulated theta (4-12 Hz) activity—or in short dLFP_θ_ (amplitude or phase or both) (Agarwal et al., 2014) and the instantaneous amplitude at ultra-high frequency (>300 Hz) band, or FPA_uhf_ ---a proxy of the unclustered multiunit activity (Stark and Abeles, 2007; Bansal et al., 2012; Smith et al., 2013), recorded from multiple recording sites could be used to reliably decode animal’s position (**Figure 1b, c**). Augmenting multiple features, such as combining clustered spike, dLFP_θ_ or FPA_uhf_ with their own activity history (**Figure 1**) or combining FPA_uhf_ with dLFP_θ_ (see **Figure 1 — figure supplement 1b**), further improved the decoding accuracy (**Figure 1c**; two-sample Kolmogorov-Smirnov test, *p*=1.4×10^−4^, 4.6×10^-5^, 1.1 ×10^-2^ for three respective panels). We systematically assessed the decoding performance using instantaneous amplitude or phase features at various frequency bands and temporal bin sizes and found that >300 Hz amplitude was most effective. In temporal windows of <300 ms the various features and their combinations were comparable, whereas at longer time intervals decoding errors increased more rapidly for LFP_θ_ than for spike-containing combinations (see **Figure 1 — figure supplement 2**). Unless stated otherwise, we used 100 ms temporal bin size, 2 cm position bin size and optimal linear estimation (OLE) decoding method for all supervised decoding analyses (see Methods section).

**Figure 1.**
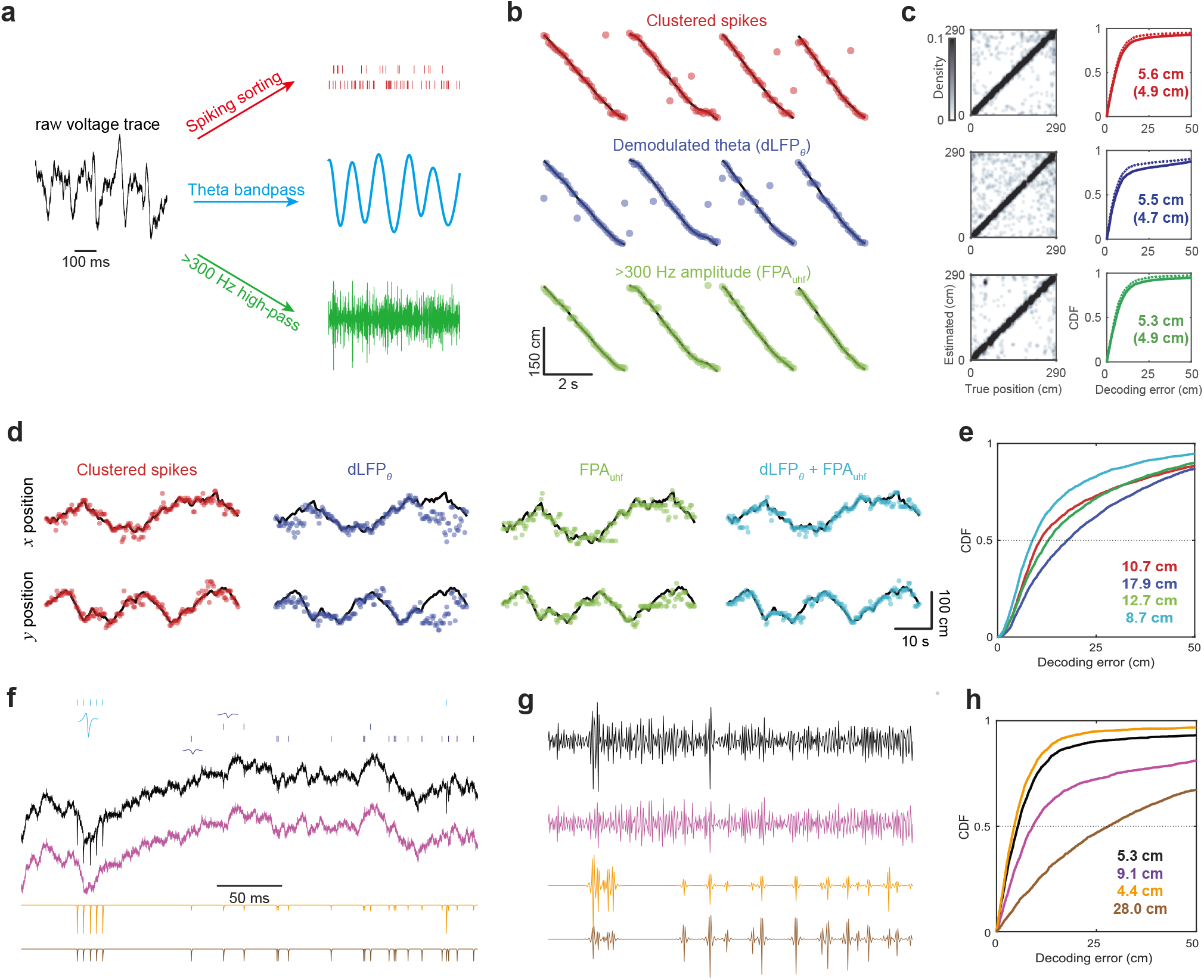
Spatiotemporal features of the hippocampus encode rat position during maze running. **(a)** Illustration of processing raw voltage traces of hippocampal recordings into three different signals: clustered spikes (red), filtered LFP at theta band (8 Hz,blue), and high-pass filtered signal (>300 Hz, green). **(b)** Decoded rat’s trajectory on a circular track (Dataset 1) based on three features derived from **(a)**: clustered spikes (red), demodulated LFP theta (referred to as dLFP_θ_, blue), >300 Hz LFP amplitude (referred to as FPA_uhf_, green). The rat’s position is marked by the black line and decoded position shown as scatter plot. **(c)** 2D histograms (left) and error cumulative distribution function (CDF; right) curves of cross-validated decoding errors during maze run (numbers indicate the median error). Augmenting the covariate with their respective history further improved the decoding accuracy in each case by ~10% (dotted line in the CDF plot, numbers in the bracket indicate the median error). **(d)** Illustration of position decoding in 2D open field environment (Dataset 5) based on the same decoding strategy derived from clustered spikes, dLFP_θ_, FPA_uhf_, and joint (dLFP_θ_+ FPA_uhf_) features. **(e)** CDFs of decoding error derived from the four different signals shown in **(d)**. Numbers indicate the median error. **(f)** Decomposition of a broadband raw voltage trace (black) into the sum of a spike-free component (magenta) and a spike-only component (orange). To investigate the effect of spike waveform and amplitude, we normalized individual spike waveforms by their respective peak amplitudes (brown). Ticks in the top represent identified spikes from clustered units and their averaged spike waveforms. **(g)** Down-sampled (to 1.25k/s) and high-pass filtered (>300 Hz) version of the four signals in (f). **(h)** CDFs of decoding error based on amplitude features derived from four signals shown in **(g)**. Number indicates the median decoding error. The following table/figure supplements are available for figure 1: **Table supplement 1.** Summary of experimental datasets under investigation. **Table supplement 2.** Summary of clustered spike-based and LFP-based decoding methods. **Figure supplement 1**. Position decoding summary from different dataset with different maze and silicon probe configurations. **Figure supplement 2**. Comparison of position decoding with different features, parameters and decoding methods. **Figure supplement 3**. Position decoding using recording sites in different hippocampal layers. **Figure supplement 4**. Comparison of position decoding between methods using standard least-squared and sparse VB-ARD regression.

Spatial coverage of the hippocampal place fields is crucial to the decoding accuracy in a larger environment. In the open field, we found that 64-channel joint dLFP_θ_+FPA_uhf_ features yielded better decoding accuracy than clustered spike-based decoding (Dataset 5; mean±SD of bootstrapped median error: 10.81±0.25 cm for spikes and 8.85±0.21 cm for dLFP_θ_+FPA_uhf_; rank-sum test, *p*=1.95×10^−21^; **Figure 1d, e**; see also **video 1**).

It is noted that the >300 Hz signal also contains spike activity (Buzsáki et al., 2012; Zanos et al., 2011; Waldert et al., 2013). To remove the contamination of spike activity, we decomposed the broadband signal into spike-free (by removing detected spikes) and spike-only components (by retaining only detected spikes; **Figure 1f**). Following down-sampling (1,250 Hz) and high-pass filtering (>300 Hz) operations (**Figure 1g**), and repeating decoding analyses showed that spike-only component produced the best decoding accuracy (**Figure 1h**), suggesting that the position decoding contribution of FPA_uhf_ was mainly derived from the spiking-related activity, although decoding the animal’s position does not require a preprocessed clustering step. To further investigate the effect of spike waveform and amplitude, we normalized individual spike waveforms by their respective peak amplitudes and did the same decoding analysis **(Figure 1f**). The results show that, in our decoding strategy, the most important information come from spike amplitude feature, which is not surprising considering our high-pass filtering and Hilbert transforming operation (**Figure 1h**). It is noteworthy to point out that even after the removal of spike waveforms, the remaining spike-free high frequency component can still yield decent decoding performance (**Figure 1h**, median error 9.1 cm derived from the spike-free component and 5.3 cm from the raw LFP).

Since spiking activity is derived mainly from the somatic layer, we further examined the impact of anatomical location of recording electrodes on decoding performance (also **Figure 1 — figure supplement 3**; Dataset 3). As expected, recording sites in the CA1 and CA3 pyramidal layers contributed most to overall FPA_uhf_ decoding accuracy. This was expected since these two layers are known to have higher cell density.

### Hippocampal FPA_uhf_ features decode replay events

During immobility and NREM sleep, putative hippocampal memory replay events occur during SPW-Rs, where population firing bursts coincide with a high ripple band amplitude (**Figure 2a**). A central task of statistical analysis is to identify the content of these replays and assess their significance in the absence of behavioral ground truth. Here we used three different decoding approaches (clustered spiked with traditional Bayesian replay decoder, clustered spiked with OLE decoder and FPA_uhf_ with OLE decoder, see Methods section) and found that all three approaches could reliably reconstruct the replay trajectories (**Figure 2b** and **Figure 2 — figure supplement 5a**). In addition, FPA_uhf_ and clustered spike-based decoding strategies produced slightly different replay trajectories (**Figure 2c**) and statistics for significant memory replay events (**Figure 2d** and **Figure 2 — figure supplement 5b**). This is interesting since all three different approaches could produce excellent decoding results in maze run state but different relay trajectories in the NREM sleep state. This suggests that the selection of feature and decoder may affect the ripple replay analysis, which raise on traditional ripple replay analysis methods. Although we have not provided a conclusive answer from our current data, our proposed approach provides an alternative means for future study of relative questions.

**Figure 2.**
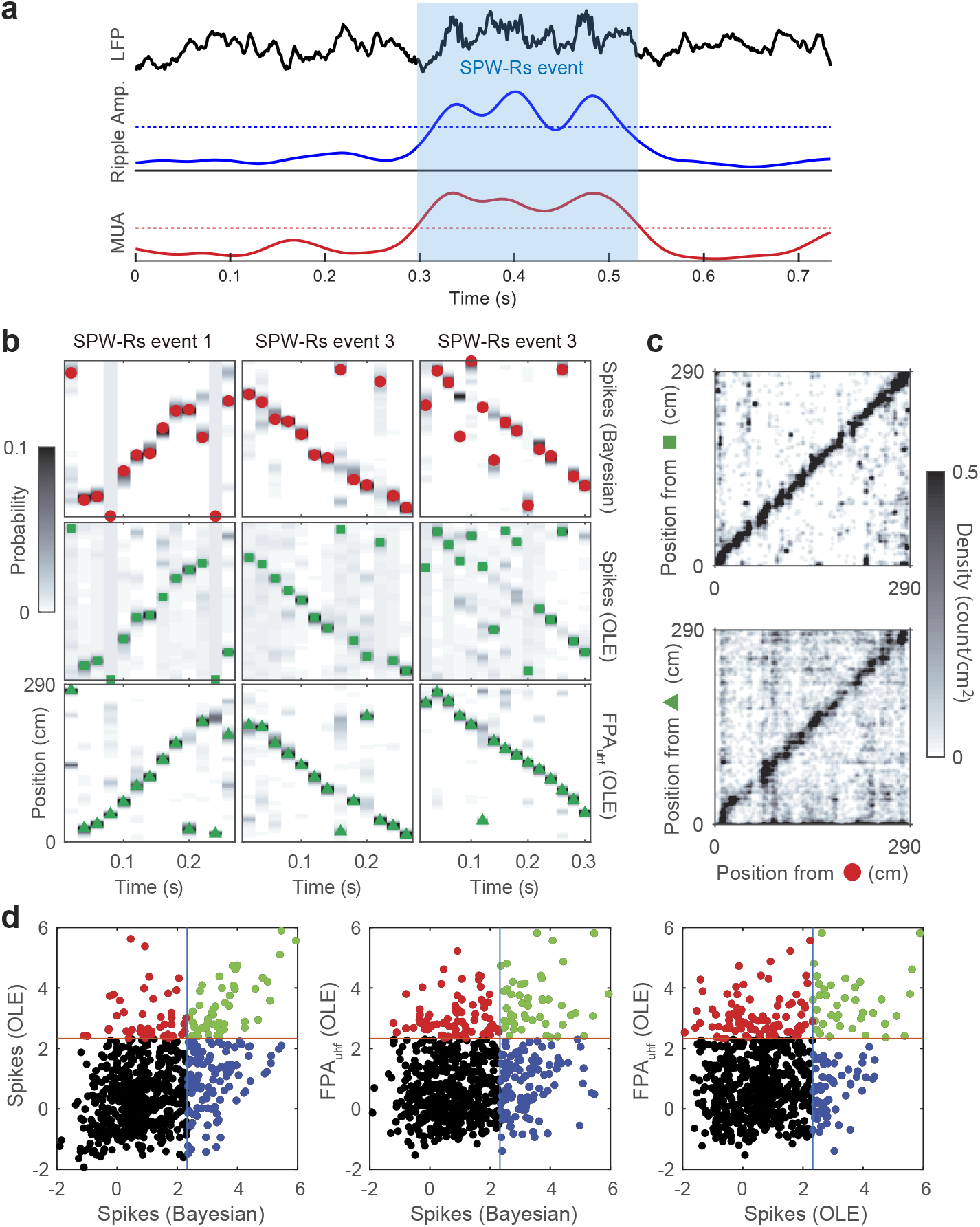
Spatial distributed hippocampal LFP patterns encode rat position during ripple events. (**a**) Illustration of a broadband hippocampal LFP (black) during a candidate replay event. The candidate event was determined by a joint criterion (see Methods section) using the ripple-band amplitude (blue) and total unclustered spike count (red, FPA_uhf_). The red and blue horizontal dotted lines indicate their respective thresholds. **(b)** Three representative example trajectories of virtual maze positions during hippocampal SPW-Rs events, derived from a traditional Bayesian decoder using clustered spikes (*top*), from an optimal linear estimator (OLE) using clustered spikes (*middle*), and from an OLE using FPA_uhf_ (*bottom*). Dark pixels show high probability values. Red circle, green rectangle and green triangle mark the estimated replay trajectories using different approaches. **(c)** 2D histogram comparisons between trajectories from three estimators shown in (**b**). **(d)** Comparison of Z-scored statistics of distance correlation (see Methods section) for ripple replay candidate events (n=727 ripples during waking and NREM sleep). Decoding methods (OLE or Bayesian) and features (FPA_uhf_ or clustered spikes) are labeled in each paired comparison. Dashed lines mark the Z-score value of 2.33 (*p*=0.01). The following figure supplement is available for figure 2: **Figure supplement 5.** Examples of decoded ripple-replay contents derived from 2nd dataset.

### Real-time decoding strategy for large-scale hippocampal recordings

Our results directly point to an efficient hippocampal FPA_uhf_ or dLFP_θ_-based decoding strategy with two key advantages: low-sampling rate requirement, easy implementation, as well as the stability and longevity of signals (in contrast to unreliable spike detection and sorting of unstable units). First, we validated the robustness of FPA_uhf_ or dLFP_θ_-based decoding strategy. In multiple consecutive recording sessions (separated by days in between) from the same animal and the same environment (Dataset 6), we trained the linear decoder using one run session and tested it on remaining run sessions. Remarkably, the FPA_uhf_ and dLFP_θ_ decoding strategies produced outstanding performance across several consecutive run sessions/days (**Figure 3a**). Furthermore, we assessed the average cross-session decoding performance when training and testing sessions were separated by different intervals, and found a V-shaped performance drop with increasing time intervals (**Figure 3b**). When the intervals between training and testing session was short, FPA_uhf_ features produced better decoding performance; yet the performance of LFP_θ_ features decayed slower. While FPA_uhf_ and dLFP_θ_ features were complementary, combining them further improved the decoding accuracy. The slowly degraded decoding accuracy in these features across days might be induced by neuroplasticity and place cell remapping (Linderman et al., 2016).

**Figure 3.**
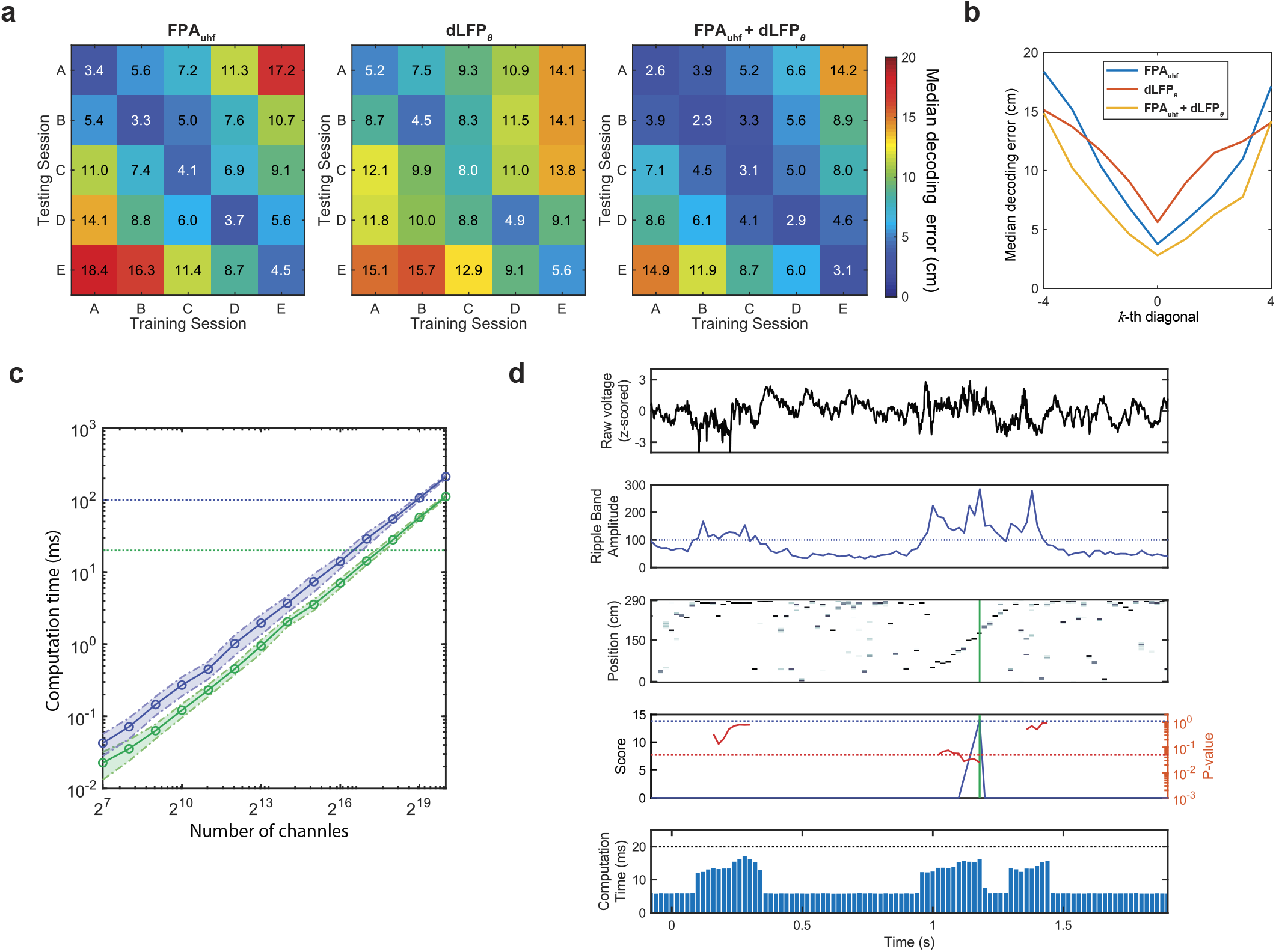
Robustness and scalability of different decoding strategies. **(a)** Cross-session decoding error matrices summarize within-session (cross-validated) and between-session assessment using dLFP_θ_, FPA_uhf_ and dLFP_θ_ + FPA_uhf_ features in five consecutive run sessions (Dataset 6, Mouse 1). The number in the matrix shows the median decoding error. Sessions (labeled by A,B,C,…E) were separated by ~24 hours. **(b)** Average statistics of the *k*-th (−4 ≤ *k* ≥ 4) diagonal of cross-session decoding error matrices in (**a**). *k*=0 implies within-session cross-validated decoding; *k*≤0 (lower diagonal) implies training on historical sessions and testing the subsequent sessions in time; *k*>0 (upper diagonal) implies the reverse order. (**c**) OLE decoding using FPA_uhf_ (green: 20 ms; blue: 100 ms bin size) with respect to the number of channels, *N*. In our computer simulations, we assumed a linear mapping with dimensionality of *N*×145 (290cm track, 2cm position bin size). Signals were down-sampled (to 1.25k/s) and high-pass filtered (>300 Hz) prior to computation. Shaded areas represent SD from 1,000 realizations. **(d)** Illustration of online detection and assessment of hippocampal memory replay during NREM sleep. First, from the raw extracellular voltage trace (1st row), the onset of candidate event was identified using a threshold criterion based on the ripple band amplitude (2nd row) from a pre-selected channel. Next, a “spatial trajectory” was reconstructed from hippocampal FPA_uhf_ (20 ms bin size; 3rd row). Once the duration of candidate event was more than 3 bins, the significance of decoded “spatial trajectory” was assessed based on online shuffling statistics (4th row), reporting the Monte Carlo *P* value (red). An accumulative score (blue) was continuously updated if *P*<0.05 (red horizontal dashed line). A significant replay was determined when the accumulative score was above a threshold (blue horizontal dashed line). The accumulative score was set to 0 at the detection onset and reset to 0 when the cumulative score threshold was reached. The computation time for online evaluation is shown in the last row. See Methods for detailed descriptions. The following table supplement is available for figure 3: **Table supplement 3.** Result comparison between online and offline significance assessment of rat hippocampal memory replay events.

Furthermore, we ran computer simulations to test the scalability of our decoding algorithm. Since the training and decoding of our algorithms are fully automatic, we could update our decoder at any time. For the sake of simplicity, we choose to use FPA_uhf_ features alone to achieve best decoding accuracy and to reduce computational complexity. Running an optimized C/C++ code on a multi-core desktop computer, the readout could accommodate a real-time speed at a scale up to hundreds of thousands of (>100,000) recording channels during both wake and sleep (**Figure 3c,** 100 ms temporal bin size for wake and 20ms for ripple replay). Next, it is important to assess the statistical significance of online decoded replay events during NREM state. We adapted a real-time spike-based replay assessment strategy (Hu et al., 2018) to accommodate FPA_uhf_ decoding (see Methods section). By presetting the pseudorandom shuffle operations (n=2,000 shuffles), we could accommodate a real-time speed (<20 ms) for both decoding and significance assessment for 128 channels recoding with 1.25k/s sampling rate in a 1D environment (**Figure 3d**), with comparable results derived from off-line significance assessment (see **Figure 3 — table supplement 3**). In addition, all these real-time scalability tests were performed on a regular desktop computer with CPU only (see Method section), the state-of-art clusterless online decoding algorithm (Hu et al., 2018) have to employ GPU hardware and special software which may increase the complexity and compromise the reliability in practical applications.

### Unsupervised learning reveals consistent representations between low-rank structures of hippocampal FPA_uhf_ with spikes of neuronal ensembles

Establishing a mapping between animal’s position and neural activity requires training samples during maze run. However, from an internal brain observer perspective, it is important for downstream structures of the hippocampus to quickly infer spatial representations without direct behavioral measures. Here we employed two latent variable models and unsupervised learning methods, nonnegative matrix factorization (NMF) and reconstruction independent component analysis (RICA), to extract lower-rank features from multi-site recorded hippocampal FPA_uhf_ (see Methods section and **Figure 4a, b**). Position-averaging on the feature matrix derived from NMF or RICA revealed localized structures in the latent state space. From the lower-rank FPA_uhf_ features (derived from NMF or RICA, followed by feature resampling; see Methods section and **Figure 4 — figure supplement 6**), we further trained an unsupervised Bayesian hidden Markov model (HMM) (Frank et al., 2004) and inferred the latent state trajectories (**Figure 4c**). During run, the lower-rank FPA_uhf_-inferred state trajectories matched well with the animal’s position as well as spike-inferred state trajectories (**Figure 4d**). During ripples, we also observed similar correspondence between two latent state trajectories derived independently from lower-rank FPA_uhf_ features and clustered spikes (**Figure 4e**). In both maze run and ripple events, the preprocessed lower-rank feature extraction was critical for learning the latent state trajectories; direct use of LFP or FPA_uhf_ features alone yield poor performance.

**Figure 4.**
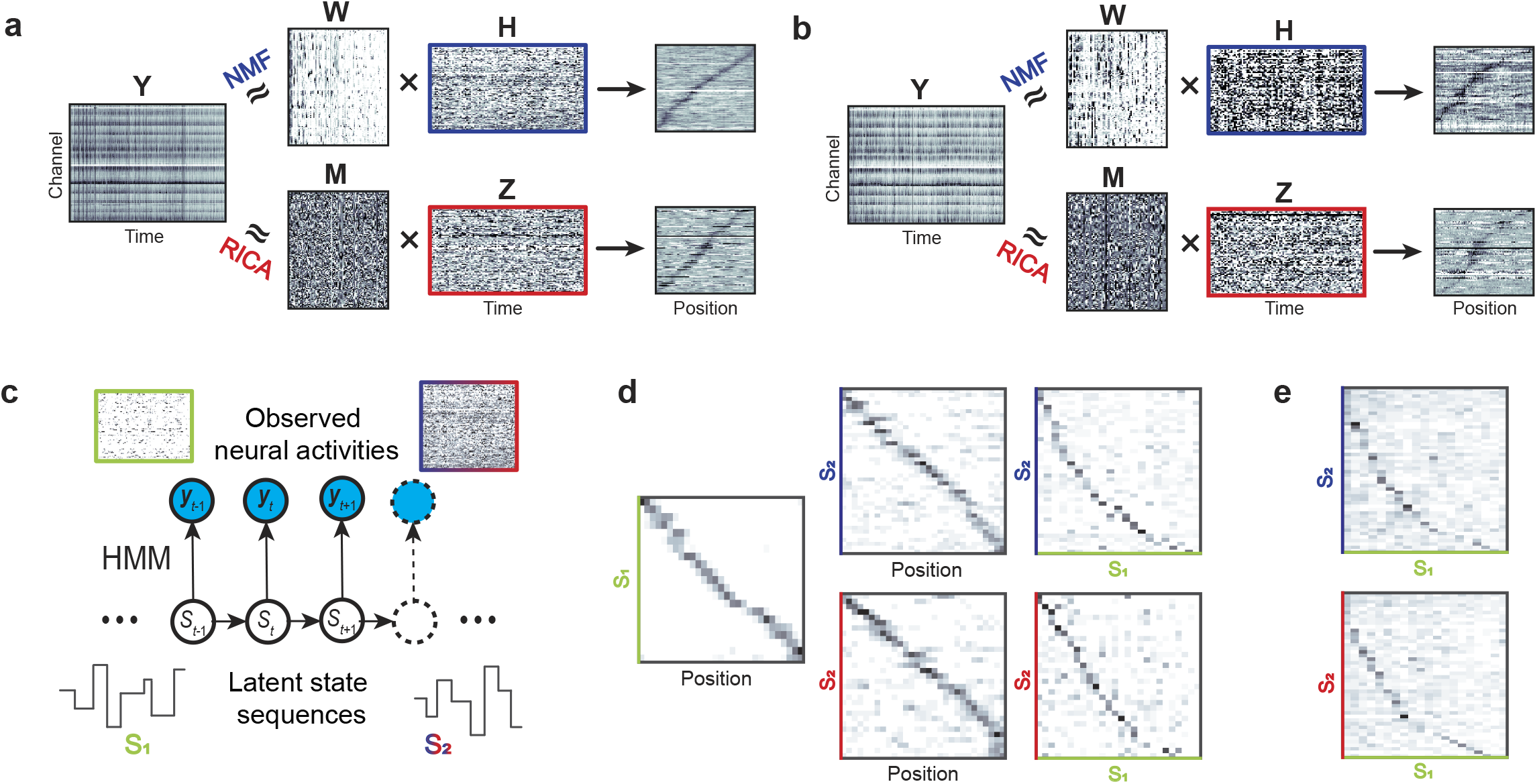
Unsupervised learning methods for extracting multichannel FPA_uhf_ features. **(a)** Illustration of NMF (*top*) and RICA (*bottom*) methods for extracting lower-rank features from FPA_uhf_ during maze run (Dataset 1). After sorting, the position-average patterns (from the rows of NMF’s matrix **H** or RICA’s matrix **Z**) yielded localized position-tuned features covering the entire track (similar to “place fields”). **(b)** Same as (**a**), during ripple events. The virtual position was decoded from clustered spikes. **(c)** A Bayesian HMM was used for inferring latent state sequences {*S*_1_(*t*)} and {*S*_2_(*t*)}, based on the spike ensembles and FPA_uhf_ features, respectively. The sorted ensemble spikes are assumed to be Poisson distributed. The lower-rank FPA_uhf_ features were obtained from the NMF’s matrix **H**, followed by a feature resampling procedure (see Methods section). The matrix frame color (green, blue, red) represent the source of feature matrix (clustered spike count, **H, Z**). **(d)** During maze running, the Bayesian HMM analysis revealed a strong one-to-one correspondence map between the animal’s position and the inferred latent states {*S*_1_(*t*), *S*_2_(*S*_2_)}, as well as between {*S*_1_(*t*)} and {*S*_2_(*t*)} (upon sorting). These consistency results indicated that the most valued information in FPA_uhf_ is spike activities (*top row*: bases on NMF; *bottom row*: based on RICA). Label color (*S*_1_ in green and *S*_2_ in blue or red) represents the latent state sequence derived from clustered spikes and lower-rank features (**H** or **Z**). **(e)** During ripples in the absence of behavior measures, the Bayesian HMM analysis also revealed a correspondence map between {*S*_1_(*t*)} and {*S*_2_(*t*)}, which were derived from the clustered spikes and FPA_uhf_ features (*top row*: bases on NMF; *bottom row*: based on RICA), respectively. Label color same as (**d**). The following figure supplements are available for figure 4: **Figure supplement 6**. Schematic of resampling procedure. **Figure supplement 7.** Dimensionality reduction of multi-site recorded FPA_uhf_ features during track running.

Furthermore, projecting high-dimensional FPA_uhf_ features onto a 2D embedding space revealed coherent structures in agreement with the animal’s position or spatial topology of the environment (**Figure 4 — figure supplement 7**). Together, these results confirmed that unsupervised learning methods can uncover neural representations of large-scale hippocampal field potentials without *a priori* measurement of animal’s behavior, which has been a challenge for inferring memory replays of hippocampal nonspatial representations or hippocampal-neocortical representations during sleep (Allen et al., 2016; Chen et al. 2016; Chen and Wilson, 2017; Maboudi et al., 2018).

### Independent, parallel and complementary hippocampal spatiotemporal patterns for spatial representation

Propagating waves in the rat hippocampus have been reported at the theta and ripple frequency bands (Agarwal et al., 2014; Lubenov and Siapas, 2009; Patel et al., 2012; Patel et al., 2013). In many cases, the spatiotemporal patterns are difficult to detected because of the lack of regular patterns. In order to fully exploit spatially distributed features of multi-site recoding, here we introduced a novel method, motivated from optical flow used in computer vision, to extract spatiotemporal patterns (see Method section), and investigated how these spatiotemporal patterns across multi-site hippocampal recordings were coordinated in space and time. We mapped the bandpass-filtered hippocampal field potentials (at 4-12 Hz or >300 Hz band) onto the 2D electrode space in time to obtain a video sequence, and then employed an optical flow estimation approach to compute the vector field between two consecutive image frames (see Methods section and **Figure 5a**). The spatiotemporally local optical flow revealed position-tuned wave patterns in the FPA_uhf_ or other LFP-derived features. Therefore, the optical flow method can help reveal the intrinsic spatiotemporal patterns of band-limited signals (beyond theta and ultra-high frequency bands). In principle, this method can also be used for other multi-site array recordings such as EEG, MEG, and ECoG.

**Figure 5.**
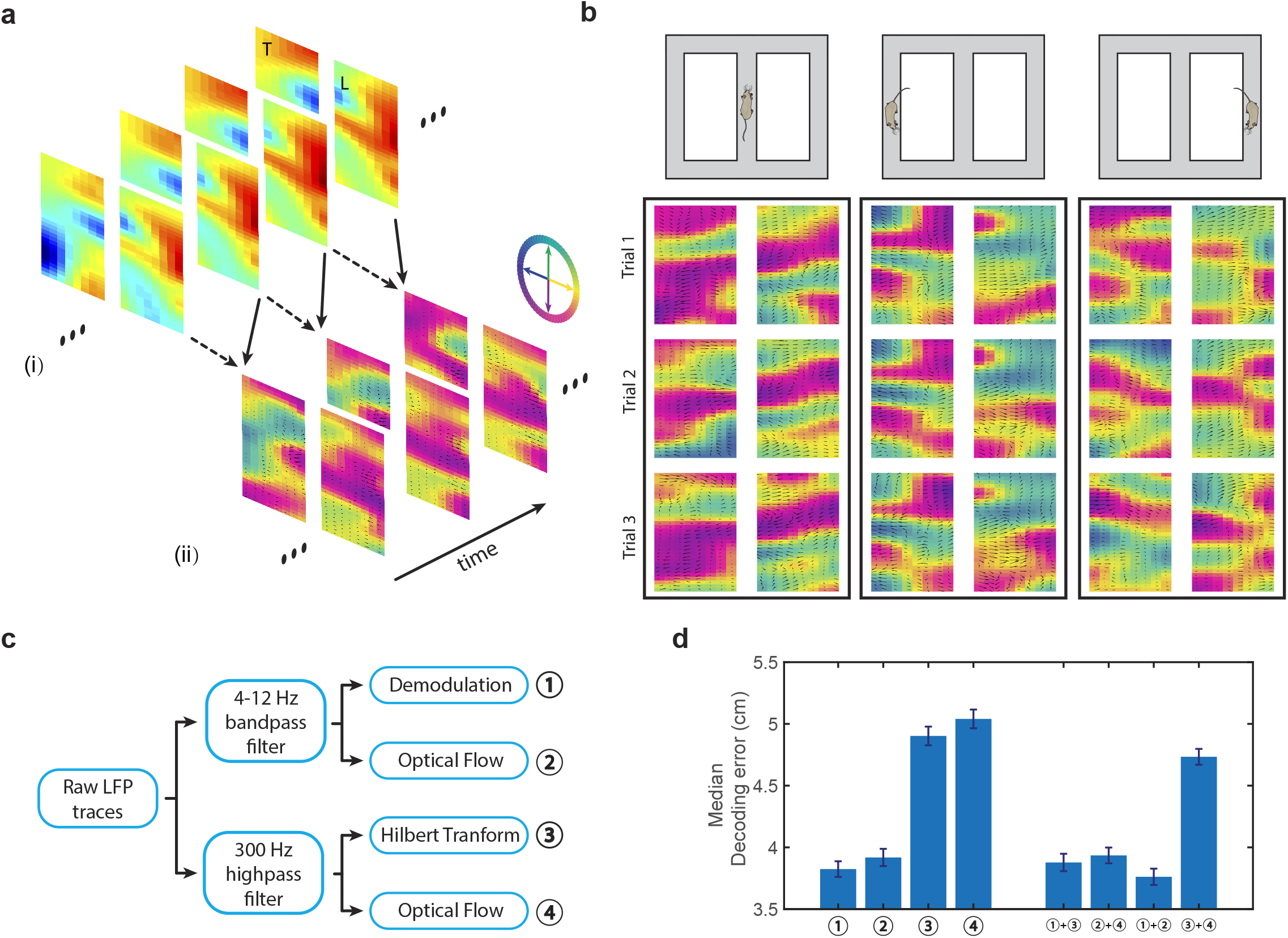
Spatiotemporal patterns of multisite recorded hippocampal LFP encode run trajectories. **(a)** Illustration of optical flow estimation and visualization by 2D vector field. High-density silicon probe recordings (8 shank, 32 sites each shank) in the same hippocampus in the transversal (T) and longitudinal (L) axes of a rat are shown separately (Dataset 3). Each 2D vector was visualized as an arrow (characterized by size and direction). (i) Background color in the raw map represents the voltage of filtered LFP traces. (ii) Background was color coded according to the arrow direction. **(b)** Optical flow estimated from band-pass filtered (4-12 Hz) 2×256-site recorded LFP signals show position-dependent patterns on a T-maze (Dataset 3). The vector field patterns showed trial-by-trial consistency at same position on maze. See Video 2 for the demonstration of clear propagating wave patterns. **(c)** Schematic of extracting parallel and independent spatiotemporal features from hippocampal field potentials at theta (4-12 Hz) and ultra-high (>300 Hz) frequency bands. Features are labeled as ①②③④. **(d)** Comparison of median position decoding error during maze run derived from four different spatiotemporal features in (**c**) and their feature combinations (Dataset 3). Error bar shows the bootstrapped SD. The following figure supplement are available for figure 5: **Figure supplement 8**. Optical flow estimated from FPA_uhf_ features and its position decoding results.

In addition to exploratory data visualization, we further investigated whether the high-dimensional optical flow features were useful for decoding. Remarkably, these spatiotemporal patterns estimated directly from bandpass-filtered hippocampal field potentials provided sufficient information for reconstructing animal’s position (LFP_θ_-flow in **Figure 5b**; see also Methods section and **Figure 5 — figure supplement 8** and **video 2**). In fact, we discovered that a set of hippocampal field potential-derived spatiotemporal patterns (**Figure 5c**) contained complementary information for position decoding (**Figure 5d** for Dataset 3; see also **Supplementary Figure 7c** for Dataset 1). The variability in contributions of individual spatiotemporal patterns in different datasets might be ascribed to spatial sampling or different layout in the implanted probes (**Figure 4 — figure supplement 1c**). Together, these hippocampal spatiotemporal features or their combinations provide a rich repertoire for position decoding across brain states (**Table 1**).

**Table 1.**
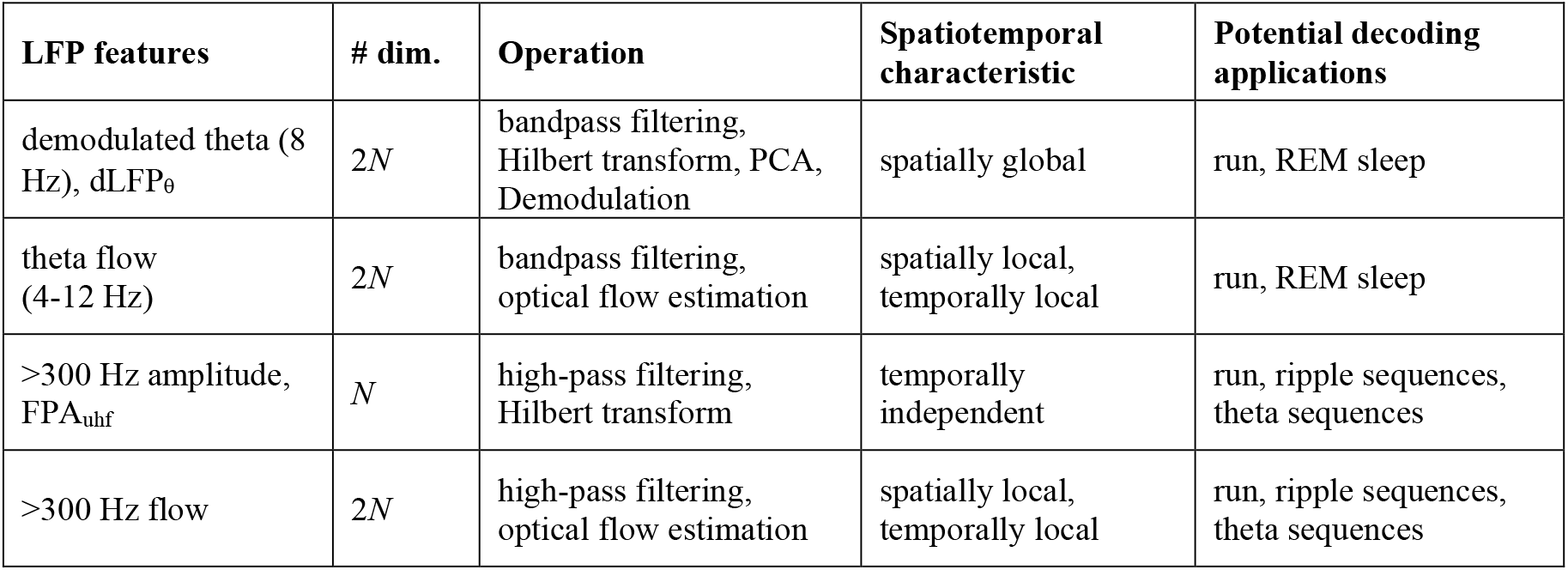
Comparison of four hippocampal LFP spatiotemporal features (*N* denotes the channel count) and their potential decoding applications.

| LFP features | # dim. | Operation | Spatiotemporal characteristic | Potential decoding applications |
| --- | --- | --- | --- | --- |
| demodulated theta (8 Hz), dLFP<sub>θ</sub> | $2N$ | bandpass filtering, Hilbert transform, PCA, Demodulation | spatially global | run, REM sleep |
| theta flow (4-12 Hz) | $2N$ | bandpass filtering, optical flow estimation | spatially local, temporally local | run, REM sleep |
| >300 Hz amplitude, FPA<sub>uhf</sub> | $N$ | high-pass filtering, Hilbert transform | temporally independent | run, ripple sequences, theta sequences |
| >300 Hz flow | $2N$ | high-pass filtering, optical flow estimation | spatially local, temporally local | run, ripple sequences, theta sequences |

### Prospective decoding strategy for predicting goal-directed navigation behavior

Hippocampal place cells exhibit trajectory-dependent or prospective/retrospective memory coding in spatial navigation (Frank et al., 2000; Ferbinteanu and Shapiro, 2003; Ji and Wilson, 2008). Here we investigated if a prospective decoding strategy based on hippocampal field potentials could predict animal’s decision in a spatial alternating task (Dataset 6). We adapted the standard decoding strategy and attempted to predict animal’s prospective L/R arm position while the animal ran in the central arm towards the choice point, with forward time lag ranging from 0.5 s to 1.5 s (5-15 temporal bins, 100 ms bin size; **Figure 6a**). Remarkably, both hippocampal spikes and joint (dLFP_θ_+FPA_uhf_) features (**Figure 6b** and see also **Figure 6 — figure supplement 9**) carried predictive representations of upcoming maze locations in the L/R arm. We found that the prediction accuracy in prospective coding improved as animals moved closer to the choice point, where the predictability was far beyond the chance level (see shuffled statistics **Figure 6 — figure supplement 9**). In two tested animals, the joint features achieved a high (86%-97%) prediction peak accuracy in correct trials (n=4 sessions for each mouse; see **Figure 6 — figure supplement 9** for incorrect trials).

**Figure 6.**
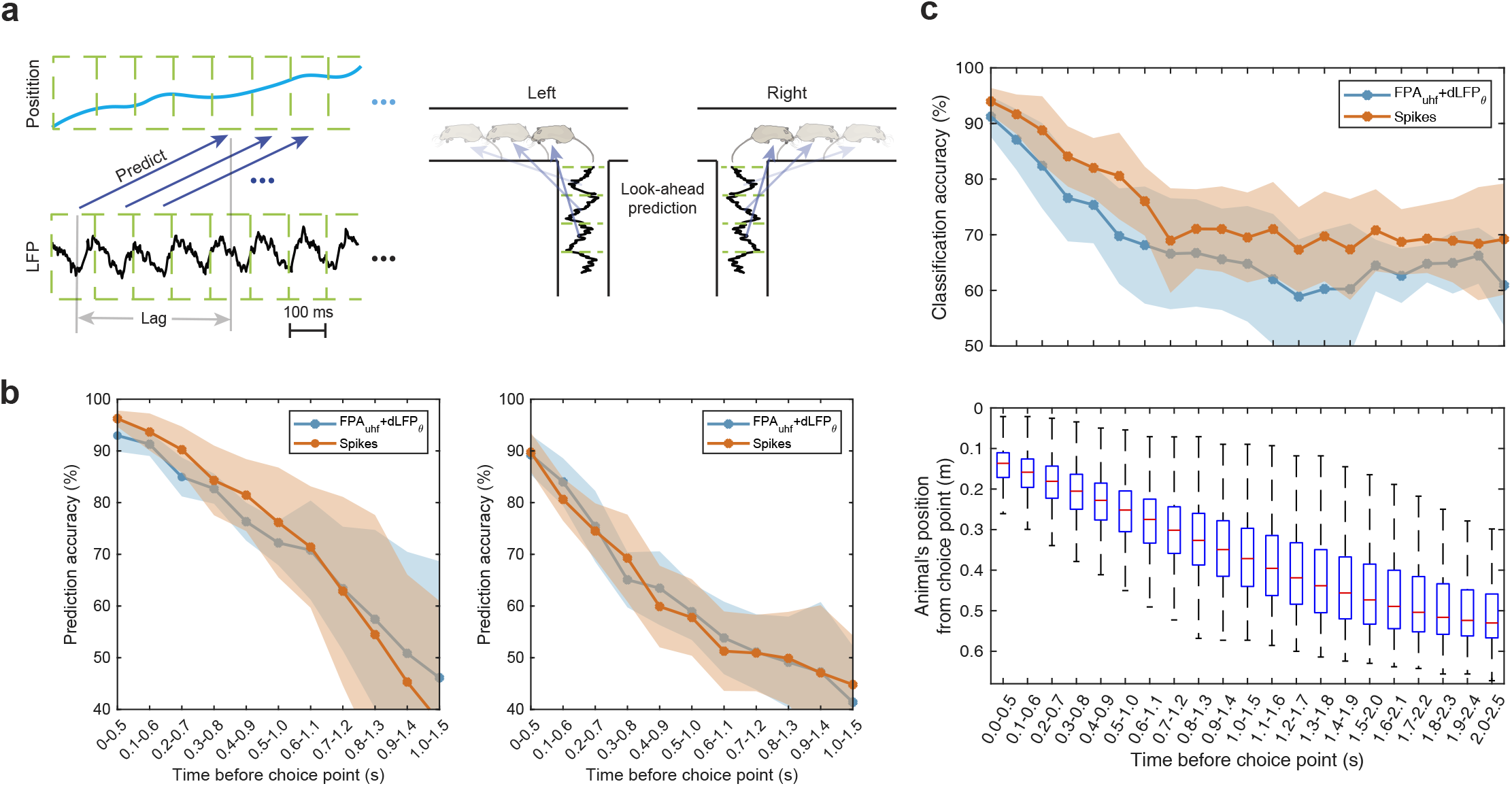
LFP, FPA_uhf_ and spike-based features predict mice’s decision-making. **(a)** Schematic of prospective decoding, with time lag of 0.5-1.5 s (5-15 temporal bins, 100 ms bin size) prior to the choice point in the T maze. **(b)** Comparison of cross-validated prediction accuracy between clustered spike-based and joint (dLFP_θ_ + FPA_uhf_) decoding strategies (Dataset 6). Mean and SD results were pooled over multiple sessions. *Left panel*: mouse 1 (n=4 sessions, n=120 correct trials per session); *right panel*: mouse 2 (n=4 sessions, n=120 correct trials per session). **(c)** *Top panel*: cross-validated SVM classification accuracy (for mouse 1). The peak classification rate from cross-validation was ~90% and the AUROC statistic was ~0.94. *Bottom panel*: Boxplot statistics of animal’s actual location at specific temporal bins across all tested trials. The following figure supplements are available for figure 6: **Figure supplement 9**. Comparison of discrimination accuracy in prospective position decoding derived from different features. **Figure supplement 10. Comparison of linear SVM classification accuracy derived from different features.Video legends**

In another independent test, we further trained a linear support vector machine (SVM) classifier to predict the L/R decision (for correct trials only) by pooling dLFP_θ_ and FPA_uhf_ features at consecutive 5 temporal bins prior to the choice point (see Methods section and **Figure 6c**). The cross-validation classification accuracy and trend was also comparable to **Figure 6b** in two tested animals (peak accuracy, correct trials: 86-98% for spikes, 77-95% for joint (FPA_uhf_+dLFP_θ_) features; incorrect trials: 66-94% for spikes, 64-89% for joint features). The gradual decay in classification accuracy away from the choice point was partially due to the increased trial variability in the animal’s actual position because of the variability in run speed. Notably, the degraded accuracy trends are slightly different between forward and backward directions, when triggering temporal bins from either the end (‘backward’) or start (‘forward’) position of the central arm (see **Figure 6 — figure supplement 10**).

## Discussion

Rodent hippocampal clustered spikes and spatially distributed theta waves are known to encode animal’s position. Here, we further demonstrated that hippocampal spatiotemporal patterns at different frequency bands contain rich spatial representations in wake and sleep. Based on large-scale multi-site electrophysical recordings of the rodent hippocampus, we proposed a set of spatiotemporal features of hippocampal field potentials and developed efficient statistical decoding methods for ultrafast readout of spatial representations. The decoding performance derived from each feature depends heavily on the layout of the silicon probe and the implanted location of hippocampus layer.

The instantaneous readout of multi-site spatiotemporal patterns of hippocampal field potentials provides a plausible mechanism of information integration along the septotemporal axis of the hippocampus, forming the traveling waves (Lubenov and Siapas, 2009; Patel et al., 2012; Patel et al., 2013). Such traveling waves can also propagate the coded information to the downstream structures of the hippocampus. Our results on FPA_uhf_ decoding were also consistent with our previous finding that unsorted hippocampal ensemble spikes can reliably encode rodent’s position (Kloosterman et al., 2014; Hu et al., 2018; Deng et al., 2015). We have also demonstrated that the prospective decoding strategy is robust and reliable to predict animal’s decision-making in goal-directed navigation. The most computationally efficient FPA_uhf_ decoding strategy is particularly useful for content-based real-time closed-loop circuit manipulation in rodents with chronic implant, which may prove critical for causal circuit dissection of hippocampal-prefrontal coordination in decision-making (Singer et al., 2013; Schmidt et al., 2019). Easy acquisition, efficient sampling, processing and transmission of band-limited signals make it ideal for wireless-empowered implanted devices.

A central task in computational neuroscience is to link neural representations with animal’s behavior or internal cognitive state. Therefore, development of unsupervised machine learning methods is important for discovering intrinsic latent structures of high-dimensional spatiotemporal neural data, especially in the absence of behavioral measures (Linderman et al., 2016; Cunningham et al., 2014; Chen et al., 2014; Townsend and Gong, 2018; Chaudhuri et al., 2019). Since high-density LFP recordings produce a high degree of input correlation between neighboring channels, low-rank feature extraction or dimensionality reduction (e.g., ICA, NMF and embedding) can help exploratory data visualization and subsequent decoding analysis.

Hippocampal spatiotemporal patterns often display in the form of traveling wave (Maboudi et al., 2018; Lubenov and Siapas, 2009; Patel et al., 2012) or sequential structure (‘hippocampal sequences’) in space and time during various behavioral state (Buzsáki and Tingley, 2018). In addition to ripple sequences, hippocampal theta sequences have been known to encode look-ahead trajectories or current goals in planning (Foster and Wilson, 2007; Gupta et al., 2012; Wikenheiser and Redish, 2015). Furthermore, internally generated sequences may predict animal’s choices in a memory task or guide navigation when external spatial cues are reduced (Pastalkova et al., 2008; Wang et al., 2015; Villette et al., 2015). To date, rodent hippocampal memory replays have been widely studied in NREM sleep. During REM sleep, rodent hippocampal LFP theta oscillations are pronounced, yet their spatiotemporal patterns are less well understood (Louie and Wilson, 2001). The spatiotemporal patterns of multi-site hippocampal field potentials and decoding methods proposed here may provide new opportunities to investigate these REM sleep-associated spatiotemporal patterns.

Although we only investigated the rodent hippocampal circuit here, our ‘place’ decoding analysis and methods can be readily applied to other neocortical areas that encode similar spatial information, such as the entorhinal cortex, primary visual cortex, retrosplenial cortex, parietal cortex and somatosensory cortex (Hu et al., 2018; Ferbinteanu and Shapiro, 2003; Hafting et al., 2005; Whitlock et al., 2008; Mao et al., 2017; Haggerty et al., 2015; Long and Zhang, 2018). Examination of the coordinated representations of large-scale hippocampal-neocortical spatiotemporal activity during various behavioral states would help dissect the mechanisms of memory, learning, planning and decision-making (Chung et al., 2019).

Finally, the superior representational power of hippocampal field potentials points to potential applications for investigating human memory replays based on high-density MEG or EEG recordings (Kurth-Nelson et al., 2016; Liu et al., 2019; Huang et al., 2018), or for predicting human hippocampal memory task outcomes based on ECoG recordings (Zhang and Jacobs, 2015).

## Materials and methods

### Spike sorting

Extracellular representations of action potentials were extracted from recorded broadband (0.3 Hz-10 kHz) signals followed by threshold-based spike detection algorithm. Individual spikes were automatically clustered using the KlustaKwik algorithm (http://klustakwik.sourceforge.net/) or Kilosort algorithm (https://github.com/cortex-lab/KiloSort). The clusters were manually refined by discarding multiunit clusters showing lack of clear refractories in the autocorrelogram, and groups with unstable firing patterns over time.

### Theta-band LFP demodulation

We applied a complex Morlet wavelet (with central frequency at 8 Hz) to the broadband LFP signal followed by (noncausal) zero-phase filtering. From the theta-band filtered analytic LFP signal, we applied demodulation using the following equation to obtain theta-demodulated LFP signals *y_d_* (*t*)

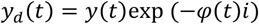

where 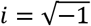, *y*(*t*) denotes the band-pass filtered signals at the theta band, *φ*(*t*) denotes the phase of the first principal component (PC) of the complex-valued, filtered multi-channel LFP signal within the theta frequency band. The first PC reflects the oscillatory component common to all channel (Agarwal et al., 2014).

### Spike removal from raw voltage signals

To remove putative spikes from the broadband signal (up to 20 kHz sampling rate), we first detect the spikes by using Kilosort algorithm. Once the spikes were identified in each channel, we removed the spikes, by using a linear interpolation from the start to end points of spike waveform or using a Bayesian spike removal algorithm (Zanos et al., 2011), and further obtained the resulting spike-free signal (see **Figure 1f**).

### Nonnegative matrix factorization (NMF)

Let **Y** denote the *N*-by-*T* matrix consisting of nonnegative *N*-channel FPA_uhf_ features. NMF is aimed at finding an approximate low-rank matrix factorization form:

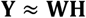

where **W** is a low-rank (*m*-by-*N, m<N*) nonnegative matrix, and **H** is an *m*-by-*T* nonnegative matrix. We interpreted the column vectors of **W** as a set of basis functions that reflected the spatially localized feature in the latent space. We varied the model order *m* and compared the visualization effects.

### Reconstruction independent component analysis (ICA)

Let **Y** denote the *N*-by-*T* matrix consisting of real or complex-valued *N*-channel LFP features at a specific frequency band. The ICA assumes that **Y** is a linear combination of independent source signals: **Y=MZ**, where **M** is a square mixing matrix, and **Z** denotes the *N*-by-*T* source signal matrix consisting of mutually statistically independent source signals. ICA is aimed to seek an optimal demixing matrix, **W**, operated on the **Y**, such that 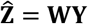 recovers the independent source signals 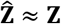. For real-valued FPA_uhf_ features, we employed a reconstruction ICA (RICA) algorithm (Le et al., 2011). In contrast to ICA, RICA replaces the orthonormal constraint with a soft reconstruction penalty and allows an overcomplete representation (i.e., **W** is a non-square matrix).

### Characterization of LFP traveling waves

To characterize the traveling wave patterns of multichannel LFP signals, we used the optical flow (vector field) method. Optical flow is commonly referred to the pattern of apparent motion of objects, and edges in a visual scene. In computer vision, the optical flow methods try to calculate the motion between two image frames which are taken at times *t* and *t* + Δ*t* at every voxel or pixel position. These methods are called differential since they are based on local Taylor series approximations of the frame images; that is, they use partial derivatives with respect to the spatial and temporal coordinates. For a 2D+*t* dimensional case, a voxel at location (*x, y, t*) with intensity *I*(*x, y, t*) will have moved by {Δ*x*, Δ*y*, Δ*t*} between the two image frames, and the following *brightness constancy constraint* can be given:

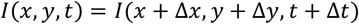

Based on first-order Taylor series expansion, we derived

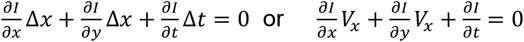

where 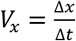 and 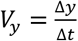. In the vector form, the above equation is:

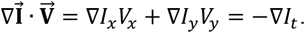

We projected the filtered (4-12 Hz or >300 Hz) multichannel LFP signals onto a 2D image according to the recording sites array layout. We then used the Horn-Schunck estimation method to estimate the optical flow based on neighboring frames (Horn and Schunck, 1981). The optical flow was represented and visualized by a 2D vector field, with arrows indicating the direction, and the size of arrow proportional to the scale. To decoding animal’s position, we further temporally averaged the vector field movie at a frame rate of 10 Hz (i.e., bin size: 100 ms). These high-dimensional flow patterns were used in a simple linear decoding method (such as OLE). In real-time applications, estimation of optical flow can be implemented via graphic processing unit (GPU), such as NVIDIA optical flow SDK (https://developer.nvidia.com/opticalflow-sdk).

### Stochastic neighbor embedding (SNE) of multichannel FPA_uhf_ features

The (*t*-distributed SNE algorithm is a robust probabilistic dimensionality reduction method for visualization of high-dimensional data (Van der Maaten and Hinton, 2008). The goal of embedding was to approximate the probability distribution of high-dimensional FPA_uhf_ features when the same operation was performed on the low-dimensional (*n*=2 or 3) “image” of the FPA_uhf_. During animal’s maze run, we color coded the embedded FPA_uhf_ features in a two-dimensional embedding space according to the animal’s linearized position.

### Decoding Analysis

Several spike-based or LFP-based decoding methods were investigated (see **Figure 1 — table supplement 2** and **Figure 1 — figure supplement 2c**). In all decoding analyses, spikes or LFP features were Z-scored across features (neurons or channels), respectively.

### Bayesian spike decoding

For decoding with sorted spikes, one method we used is traditional replay reconstruction algorithm (Zhang et al., 1998). According to Bayes’ rule, the probability of animal at position *x* given the spiking activities *y* in a short window is:

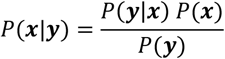

By assuming a Poisson firing for each unit, the likelihood function written as:

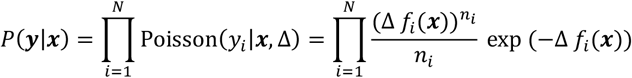

where *f_i_*(*x*) donates the average firing rate of unit *i* at position *x*, which can be inferred from the maze run data (ripple event analysis, **Figure 2**) or training data (position decoding analysis, **Figure 1d**). Δ is the temporal bin size, and *n_i_* is the spike count of unit *i* within that window Δ during the ripple event. Combining equations above yields posterior probability:

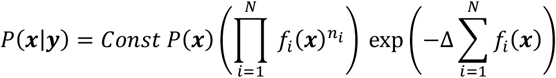

Where *Const* is a normalization factor, which can be calculated by the normalization condition ***∑_x_****P*(***x***|***y***) = 1. *P*(*x*) denote the prior probability, which can be assumed as a uniform distribution (**Figure 2**), and the Bayesian decoder reduces to a likelihood-based decoder. Alternatively, *P*(***x***) can depend on the previously decoded position^14^ (Gaussian filter, **Figure 1d**):

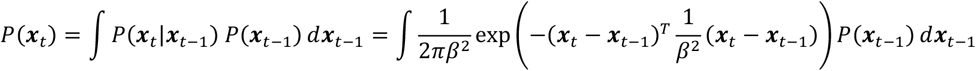

Finally the reconstructed position 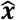 is derived from the maximum a posteriori (MAP) estimate:

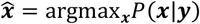

### LFP decoding based on optimal linear estimation (OLE)

We used the standard OLE method previously described before (Agarwal et al., 2014). Let *x* denote the decoded variable, and ***y*** = {*y_c_*} denote the *C*-dimensional observation. In one-dimensional environment, we mapped the linearized position *x* (with length *L*) via *K* equally spaced von Mises functions characterized by a circular variable *θ* = 2*πx/L*: *B_k_* (*θ*) = exp (*K*cos(*θ* – *θ_k_*)) for *k*=1,…,*K*. The OLE tries to solve a least-squared (LS) estimation problem:

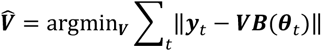

Where ***V*** = {*υ_c,k_*} denotes the unknown parameters, and ***y***_*t*_ denotes the vectors of spike count or multichannel LFP features at time *t*, and ***B***(***θ***_*t*_) = [*B*_1_(***θ***_*t*_),…., *B_K_*(***θ***_*t*_)] denotes a *K*-dimensional vector that expands the position in the basis. Unless stated otherwise, we used *K*=75, *κ* = 100 and a temporal bin size of 100 ms (maze run) or 20 ms (ripple events). In all decoding methods, all LFP features were first averaged within each time bin and then Z-scored across all channels. We rescaled the OLE output between 0 and 1 via a softmax operation.

In addition, we considered incorporating the prior history of LFP activity (such as ***y***_*t*–1_) as additional covariance in the linear regression estimator. However, this was at the cost of doubling computational complexity and potential overfitting.

In the open field, we mapped the two-dimensional position ***x*** by replacing *K* von Mises basis functions with a set of two-dimensional Gaussian tiling (Agarwal et al., 2014)

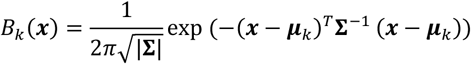

Where the vectors ***x*** represents the animal’s 2D position, and **Σ** denotes the (diagonal or isotropic) covariance, and {***μ***_*k*_} denotes the mean vectors of *K* Gaussians that tile the field. We used *K*=144 and temporal bin size of 300 ms for the open field environment.

### LFP feature likelihood-based decoding

We used a Gaussian likelihood-based decoder (i.e., noninformative prior) based on LFP features

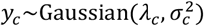

where 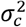 denotes the variance, and the mean *λ_c_* is represented by a sum of one or two-dimensional basis functions

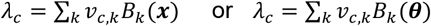

The Gaussian log-likelihood function, denoted as *£*, is written as

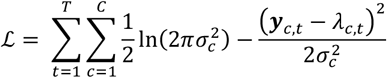

where *λ_c,t_* = ∑_*k*_ *υ_ck_B_k_*(*x_t_*). The unknown parameters 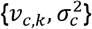 were estimated from the maximum likelihood estimation.

### OLE decoding by imposing a sparsity constraint

In the OLE method, the traditional linear least-squared (LS) regression method is subject to overfitting in the presence of high-dimensional features. To improve generalization and assist variable, we used a sparse Bayesian linear regression method based on variational Bayes (VB) and automatic relevance determination (ARD) (Drugowitsch, 2013). We extended the multi-input single-output (MISO) regression problem to a multi-input multi-output (MIMO) regression problem. Finally, the VB inference produced the posterior mean and posterior variance of the individual parameter in ***V*** = {*υ_ik_*}. For variable selection, the parameter with a small mean and a small variance would be discarded. See **Figure 1 — figure supplement 4** for an illustration of encoding and decoding application with VB-ARD.

### Bayesian hidden Markov model (HMM) for unsupervised learning analysis

To discover latent structures of large-scale hippocampal population codes, we used a HMM for analyzing hippocampal ensemble spike activity during spatial navigation and sleep (Linderman et al., 2016; Allen et al., 2016; Chen et al., 2014). In a basic HMM, we assumed that the latent state process follows a first-order discrete-state Markov chain {*S_t_*} ∈{1,2,···,*N_s_*}, and the observations of neural activity at discrete time index *t*, follow a conditional probability distribution (conditioned on the latent state *S_t_*). In the case of nonnegative features (either in the form of neuronal spike counts or multisite FPA_uhf_ features), we assumed a Poisson probability with associated tuning curve functions **Λ** = {***λ***_*c*_} = {*λ_c,i_*}. The joint probability distribution of observed and latent variables is given by

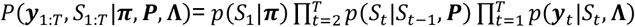

where ***P*** = {*P_ij_*} denotes an *N_s_*-by-*N_s_* state transition matrix, with *P_ij_* representing the transition probability from state *i* to *j*; ***π*** = {*π_i_*} denotes a probability vector for the initial state *S*_1_; *y_c,t_* denotes the number of spike counts from cell *c* within the *t*-th temporal bin (bin size: 100 ms during wake and 20 ms during ripples) and ***y***_1:*T*_ = {*y_c,t_*}_*C*×*T*_ denotes the time series of *C*-dimensional neural response vector. In the case of spike counts, we assumed the conditional probability distribution has a factorial form: 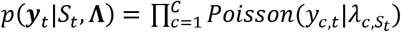, which defined the products of conditionally independent Poisson distributions. We used a Bayesian inference procedure to identify the unknown parameters {***π, P, Λ***} and the latent state sequences {*S*_1:*Γ*_} ^45^(Chen et al., 2014). We further used a Bayesian nonparametric version of the HMM, the hierarchical Dirichlet process-HMM (HDP-HMM), which extends the finite-state HMM with a nonparametric HDP prior, and inherits a great flexibility for modeling complex data (Linderman et al., 2016). The associated Markov chain Monte Carlo (MCMC)-based inference algorithm allowed us to infer the optimal model order *N_s_*.

### LFP feature resampling

In the HDP-HMM, the assumed observations were Poisson-distributed spike counts (nonnegative integers). Therefore, we need to adapt the likelihood model of the HMM to accommodate hippocampal field potential observations. In the case of nonnegative features *y_t_* (e.g., derived from NMF operated on nonnegative FPA_uhf_ or dLFP_θ_ power), we generated the same size of random samples from a desired Poisson distribution based on a rank-invariant resampling procedure (Honey et al., 2009), and replaced the original samples with the ordered new samples (by keeping their rank or order unchanged). We treated 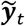 as the pseudo spike count observations and then repeated the same Bayesian inference procedure assuming a new Poisson likelihood 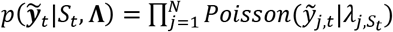. For the real-valued Z-scored features, a similar resampling procedure was also employed. A schematic illustration of the resampling procedure is shown in **Supplementary Figure 7**. The choice of the mean statistic in the new Poisson distribution was ad hoc, yet the final performance was robust with respect to a wide range of the Poisson mean statistic.

### Identification of ripples and awake and sleep replay candidate events

We first used the electromyography (EMG) and LFP for sleep staging. NREM sleep was primarily determined by the low EMG amplitude, high delta/theta power ratio, the presence of slow waves and sleep spindles in LFP activity. REM sleep was determined by the low EMG amplitude and high theta/delta power ratio. SPW-Rs were identified based on a previously described method (Grosmark and Buzsáki, 2014). The integrated power of the filtered LFP signal was calculated in a sliding window for each electrode. To identify the candidate events of memory replay during immobility and NREM sleep (see **Figure 2a**), we used a combined criterion of hippocampal LFP amplitude at ripple band (140-250 Hz) and total spike count (threshold >mean+3 SD). For visual inspection, we computed the spectrogram to assist identification. We selected subsets of candidate events during immobility (in the maze or sleep box) and post-NREM periods for FPA_uhf_ or clustered spike-based decoding analysis.

### Prospective decoding and prediction of animal’s future spatial decision-making

In goal-directed navigation tasks, we developed a look-ahead decoding strategy to predict animal’s future decision (L vs. R turns) based on either ensemble spikes, dLFP_θ_, or FPA_uhf_ features collected in the central arm of T-maze. During the training phase, we correlated the animal’s future position at the L/R arm with neural activity with a time lag (0.5-1.5 s, 100 ms temporal bin size) while animal ran in the central arm (see **Figure 6a** for schematic illustration). We assessed the “mode” of decoded L vs. R-turn trajectories and determined the prediction outcome based on their sums of weighted scores---the one with a higher cumulative sum was deemed the winner. The score at each time point was weighted with a slow decay from the future to the past. For instance, we used a linear decay weight vector [1, 0.9, 0.8, 0.7, 0.6], by imposing a larger weight on the future predicted outcome (yet the ultimate prediction outcome was insensitive to the exact values in the decay weight vector). To assess the prediction accuracy, we conducted 10-fold cross-validation. In addition, we ran 1,000 Monte Carlo runs (by randomly selecting 90% training trials in each run) and reported the mean±SD prediction accuracy. As control for each strategy, we also performed 1,000 random decoder channel order shuffles and computed the chance-level prediction accuracy.

Furthermore, we trained a linear support vector machine (SVM) classifier to predict animal’s future decision based on different features. For the ease of interpretation, we used a linear kernel in the SVM. In addition to the classification accuracy, we also reported the AUROC (area under ROC curve) statistic. An AUROC value of 0.5 indicates a chance level, whereas a high AUROC value (close to 1) implies excellent performance.

For all the prediction analyses, we only used the trials that animal start from the beginning of center arm and run directly to the next left/right arm without hesitation or stopped before the choice point. By excluding these invalid trials, we reduce the possibility that animal will change its decision throughout center arm run.

### Real-time FPA_uhf_ decoding based on the OLE method

Our real-time FPA_uhf_ decoding operation consisted of three steps. *Step 1*: Broadband voltage signals were filtered (>300 Hz) in data acquisition hardware. *Step 2*: In the memory buffer (20 ms during ripples and 100 ms during run), multichannel LFP signals were processed by a Hilbert transform in parallel (using multi-core CPU), and instantaneous LFP amplitude features were computed in raw 1.25 kHz sampling rate and then averaged. The memory was constantly updated in time. *Step 3*: Position was reconstructed based on a pretrained linear map ***V***from previous session. The pipeline was initially implemented in MATLAB R2019a (MathWorks), and optimized by C/C++ implementation (with Intel c++ compiler). We tested all real-time operations on a desktop (3.6 GHz 4-core Intel^®^ i7700 CPU, with Windows^®^10 OS, Visual Studio 2017 and Intel^®^ Parallel Studio XE 2018 environment).

Once the LFP features were computed across channels and averaged within each time bin, the remaining computational complexity of OLE operation was *O*(*N*L*) per time bin (where *N* denotes the number of channels and *L* denotes the number of spatial bins), which was split into multiple threads in parallel for computational speedup. The computational bottleneck was the Hilbert transform.

### Real-time significance assessment of decoded memory replay

To assess the significance of the FPA_uhf_ decoding result, we ran shuffle analyses. To accommodate the real-time computation, we performed and saved two types of pseudorandom shuffles (i.e., channel order shuffle by permutating columns and receptive field shuffle by circular-shifting each column) in advance and directly applied those in online decoding.

We used Hilbert transform to computed the LFP amplitude at ripple band (140-250 Hz) for a preselected channel that shows the largest ripple amplitude, and then compared ripple band amplitude with a predetermined threshold (e.g., mean+SD) to detect the onset of candidate event (**Figure 3d**). Since the decoding speed is ultrafast, we used a “decode-as-you-go” strategy. Specifically, we calculated >300 Hz high-pass filtered amplitudes features (FPA_uhf_) for all recording channels and continuously decoded the content for each time bin. But the significance assessment of the candidate event was only ran after the length of event above 3 bins. Once the significance assessment is started, we ran 2,000 (1,000 for each type of shuffle operation) random shuffle decoding analysis for the whole event at each time step and computed the Monte Carlo *P*-value (based on the distance correlation measurement). After the P-value is calculated, we update the cumulative score as follows

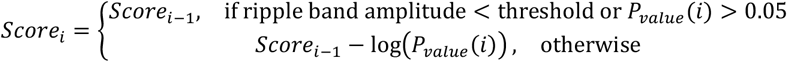

where *i* denotes the time bin index, and *Score*_0_ = 0 at the time of event onset. Once the online cumulative score was above a predetermined threshold (e.g., −3log (0.01) used in our current analyses), the event was deemed statistically significant and we reset the cumulative score to be 0.

### Statistical assessment of decoding

During maze run (velocity threshold: 5-15 cm/s depending on the spatial environment), we evaluated the decoding accuracy by the absolute error between the animal’s actual position and the decoded position: 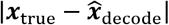 (where 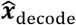 was derived from a specific decoding strategy using either clustered spikes, FPA_uhf_ or dLFP_θ_ features). We used 10-fold cross-validation to assess the median decoding error, and computed the bootstrapped standard deviation (SD).

During ripple events, we computed the distance correlation (Liu et al., 2018), and then computed 2,000 random shuffles for each event to derive the Z-score statistic or Monte Carlo *P*-value. For spike-based likelihood decoding, we used cell identity shuffling and receptive field shuffling; for FPA_uhf_ decoding, we used channel order shuffling and linear map shuffling (with respect to the rows of matrix ***V*** from OLE). Among different types of shuffling operations, we used the worst Z-score or Monte Carlo *P*-value as the final shuffle statistics.

### Region-specific assessment of decoding

To assess the region-specific contribution to specific decoding strategy, we first split the channels according to their implanted anatomical regions by using current source density map during sharp wave ripple (**Supplementary Figure 3**). To control the imbalance of channel counts between regions, we randomly selected the matched number of channels from other regions and repeated the same decoding analysis. We repeated the random selection for 1,000 times and compared their Monte Carlo statistics.

## Acknowledgments

We thank G. Agarwal and I.H. Stevenson for valuable discussions. We also thank the valuable feedback and support of G. Buzsáki and the Buzsáki lab members for sharing the experimental data. This work was partially supported by the US National Science Foundation (NSF) grant CBET-1835000 (Z.C.), NIH grants R01-NS100065 (Z.C.), R01-MH118928 (Z.C.), L.C. was supported by the China Scholar Council fellowship (CSC201806140122). V.V. was supported by a Marie Curie Actions International Fellowship (HippAchoMod, 707359).

## Competing interests

The authors declare no competing interests.

## Author contributions

Z.S.C. and L.C. conceived this research and designed the experiments; L.C. and Z.S.C. analyzed the data; V.V. performed the mouse experiments; Z.S.C. wrote the paper; and all authors participated in the editing and revisions of the manuscript.

## Additional information

Supplemental information includes ten figures, three tables, and three videos.

## Data and software availability

All data needed to evaluate the conclusions in the paper are present in the paper and/or the supplementary materials. Some raw datasets included in this study are publicly available at https://crcns.org/data-sets/hc/ or https://buzsakilab.nyumc.org/datasets/. The rest of raw data are available upon reasonable request. Some processed data and custom MATLAB/Python scripts can be downloaded from www.cn3lab.org/software.html or github repository: https://github.com/lcaopcn/Spatiotemporal-patterns-of-rodent-hippocampal-field-potentials-uncover-spatial-representations.git

---

**Video 1.** Demonstration of Bayesian (with Gaussian filter prior probability) decoding decoding in an open field environment (Dataset 5) based on clustered spikes, dLFP_θ_, FPA_*uhf*_, FP joint (dLFP_θ_+ FPA_*uhf*_) features, and combining all features. Video frame: 3.3 Hz.

**Video 2.** Demonstration of location-tuned spatiotemporal patterns in 512-channel rat hippocampal LFP signals filtered at 4-12 Hz (Dataset 3). Animal’s actual position (grey) and decoded position (blue) are shown. The 2D vector field illustrates the optical flow patterns across two Buzsáki256 probes (transversal (T) and longitudinal (L) axes in the same hippocampus). Background was color coded according to the arrow direction. Video frame rate: 10 Hz.

**Video 3.** Demonstration of location-tuned spatiotemporal patterns in 120-channel rat hippocampal FPA*uhf* features while running in a circular maze (Dataset 1). Animal’s actual position (grey) and decoded position (blue) are shown. The 2D vector field illustrates the optical flow patterns across two Buzsáki64SPL probes (6×10 layout, left and right hippocampus). Background color shows the FPA*uhf* (warm color represents high value). Video frame rate: 10 Hz.

## Notes

### Competing Interest Statement

The authors have declared no competing interest.

