## Supplemental Information for "Spatiotemporal patterns of rodent hippocampal field potentials uncover spatial representations"

### STAR METHODS

### KEY RESOURCES TABLE

| REAGENT or RESOURCE | SOURCE | IDENTIFIER |
|---|---|---|
| Deposited Data | | |
| Datasets 1, 2 | https://crcns.org/data-sets/hc/hc-11 | N/A |
| Datasets 3, 4 | http://buzsakilab.com/wp/datasets/. | N/A |
| Dataset 5 | https://crcns.org/data-sets/hc/hc-3 | N/A |
| **EXPERIMENTAL MODELS: ORGANISMS/STRAINS** | | |
| Long-Evans rats | Charles River Labs | RRID: RGD_2308852 |
| Mice C57BL/6J | The Jackson Laboratory | stock no.: 000664 |
| **SOFTWARE AND ALGORITHMS** | | |
| MATLAB | MathWorks | N/A |
| PYTHON | Open source | N/A |
| Custom code for various decoding methods | This paper | N/A |

### EXPERMIMENTAL MODEL AND SUBJECT DETAILS

All experimental studies were performed in accordance with the National Institutes of Health (NIH) *Guide for the Care and Use of Laboratory Animals* to ensure minimal animal use and discomfort, and were approved by the New York University School of Medicine (NYUSOM) Institutional Animal Care and Use Committee (IACUC). Long-Evans rats (n=3) and mice (n=2) were used in the behavior tasks, and were maintained on a 12:12-h light-dark cycle and housed individually with access to food and water.

#### Electrophysiological Recordings

***Datasets 1 and 2: Circular track and linear track (rat 1).*** Male Long-Evans rats were bilaterally implanted with two 6-shank silicon probes (128 channels in total) parallel to the septo-temporal axis of the dorsal hippocampus. The silicon probe is a custom Buzsaki64SPL probe (NeuroNexus). This 6-shank silicon probe had 10 sites at each shank, and there were 4 extra channels spaced every 1.25 mm starting 1.25 mm from the tip of Shank 4 (6×10+4=64 channels). All sites were vertically staggered along the shank with 20 μm spacing between sites (**Supplementary Figure 1c**). We selected one rat ('Achilles') in the analysis. The recording session consisted of a long (~4 hour) pre-RUN sleep epoch in a familiar room, followed by a RUN epoch (~45 minutes) in a novel circular maze (1 m diameter, Dataset 1) or a linear track (1.6 m long, Dataset 2). After the RUN epoch the animal was transferred back to its home cage in the familiar room where another long (~4 hour) post-RUN sleep was recorded. Details of experimental protocols and data have been published (**Chen** et al., 2016; **Grosmark and** Buzsáki, 2016), . The electrophysiological data are publicly available (https://crcns.org/data-sets/hc/hc-11/).

***Datasets 3 and 4: T-maze and linear track (rat 2)***. Two 256-channel (512 channels in total) custom-made silicon electrodes were implanted to the right hippocampus of the rat. The silicon probe is a custom Buzsaki256 probe with 32×8 array layout (**Supplementary Figure 1c**). The two probes were perpendicularly aligned to each other in order to record along both the septotemporal and subiculo-fimbrial axis of the hippocampus. In the T-maze, rat was trained to perform a delayed alteration task, in which the animal had to choose either the left or the right arm at the decision point. After returning to the start area, the rat was confined for 10 seconds. In the following trial, the rat had to choose the opposite direction and would obtain water reward with a correct choice. In the linear track, animal simply foraged back and forth to collect water reward. Details of experimental protocols have been published (Berényi et al., 2014), and data are available at http://buzsakilab.com/wp/datasets/.

***Dataset 5: Open field (rat 3)***. Male Long-Evans rats foraged and chased randomly dispersed drops of water or food on an elevated square platform (120 ×120 $cm^2$). The silicon probe consists of two 32-channel (4×8) array (**Figure S1C**). The probe was implanted in the rat's left and right dorsal hippocampi. Data are available at https://crcns.org/data-sets/hc/hc-3

***Dataset 6: Circular T-maze (mouse 1 and mouse 2)***. Male mice were trained to perform a continuous spatial alternation task in a circular T-maze. The silicon probe consists of a 64-channel (4×16) poly2 layout (**Supplementary Figure 1c**). Each animal's recordings consisted of multiple sessions during 4-6 consecutive days, and each session lasted 50-60 min. In each session, animal was able to run 120-160 trials, with a minimum of 120 correct trials. The probe position was not changed across sessions in order to assure the stability of LFP channels. Spikes were sorted separately for each session, yielding varying number of units (**Supplementary Table 1**).

### Supplementary figure legends

**Figure supplement 1**. Position decoding summary from different dataset with different maze and silicon probe configurations. (**a**) Schematic of four spatial environments using Datasets 1-6. (**b**) Summary of median position decoding error during maze run derived from four different features. Error bar shows the bootstrapped SD. The following default setup was used: 100 ms bin size; OLE decoding method using sparse Bayesian regression for rat 1,2 and moue 1,2; Bayesian filter decoder with a Gaussian temporal prior for rat 3. The variabilities across animals is largely part of electrode type used. $dLFP_{\theta}$ decoding was superior in rat 1 because large numbers of sites across all layers of the hippocampus was used but only a fraction of those were in the pyramidal layer. The number of clustered single units in each session are indicated above the spike bar. (**c**) Configuration of the recording silicon probes used in each dataset.

**Figure supplement 2**. Comparison of median position decoding error results derived from different band-pass filtered LFP features (phase, amplitude, phase+amplitude) at (**a**) different frequency bands, (**b**) different temporal bin size, and (**c**) different decoding methods (see **Figure 1 — table supplement 2**; Dataset 1). Shaded area or error bar shows the bootstrapped SD. (**d**) Same as (**c**), except for Dataset 3 using a different configuration of silicon probe.

**Figure supplement 3**. Position decoding using recording sites in different hippocampal layers. (**a**) Sharp wave ripple triggered current source density (CSD) analysis revealed distinct hippocampal LFP sinks and sources in different anatomical layers (256 channels, 8 shanks, Dataset 3). (**b**) $dLFP_{\theta}$ (*top*) and $FPA_{uhf}$ (*bottom*) median position decoding error using recording sites in different layers (blue) compared to median position decoding error (error bar: SD) derived from 1,000 random selections of matched number of recording sites (red). (Pyr: stratum pyramidale; Rad: stratum radiatum; Lm: stratum lacunosum moleculare; DG: dental gyrus; CA3p: CA3 pyramidal layer). The number inside parentheses indicates the channel count within that layer/region. CA1 layers with changing layer-by-layer theta phase shift gave best position decoding by $LFP_{\theta}$ decoding strategy, while $FPA_{uhf}$ was most effective in CA1 and CA3 pyramidal layer.

**Figure supplement 4**. (**a**) Comparison of median decoding error results (Dataset 1) between methods using standard least-squared (LS, blue) and sparse VB-ARD (red) regression while testing different combinations of LFP-based features (①: $dLFP_{\theta}$ amplitude and phase; ②: optical flow at 4-12 Hz band; ③: >300 Hz instantaneous amplitude; ④: optical flow at >300 Hz band ($FPA_{uhf}$). *, $P<0.05$; **, $P<0.01$; *** $P<0.001$, rank-sum test. Error bar shows the bootstrapped SD. (**b**) Comparison of the mapping $\boldsymbol{V}$ between the VB-ARD (*left*) and LS (*right*) estimators used in the OLE method ("①+②" LFP features combination). The weight coefficients learned from the VB-ARD estimator were sparse.

**Figure supplement 5**. Examples of decoded ripple-replay contents derived from 2nd dataset. (**a**) Examples of decoded ripple-replay contents derived from clustered spikes and $FPA_{uhf}$ features in post-run NREM sleep (Datasets 2). (**b**) Comparison of detection statistics of hippocampal memory replay candidates in awake and NREM sleep (n=1,038 events) derived from clustered spikes and $FPA_{uhf}$ features.

**Figure supplement 6**. Schematic procedure of resampling to rescale the Z-scored real-valued observations $y$ (from an empirical distribution) to nonnegative count observations $\hat{y}$ (from a target Poisson distribution with mean statistic of 10). The red arrows indicate the conceptual procedure. Note that the order statistics remained unchanged between two cumulative distributed function (CDF) curves.

**Figure supplement 7**. Dimensionality reduction of multi-site recorded FPA$_{uhf}$ features during track running, produced by *t*-distributed stochastic neighbor embedding (t-SNE) algorithm. Each dot was color-coded in relation to the animal's position on the track. Two multi-site high-dimensional FPA$_{uhf}$ features were similar if they were close in the two-dimensional embedding space. (**a**) For the circular track (Dataset 1), we recovered a continuous trajectory pattern in the embedding space. (**b**) For the linear track (Dataset 2), we recovered direction-specific trajectory pattern in the embedding space. (**c**) For the T-maze (Dataset 3), we recovered distinct clustered patterns, corresponding to L and R turns, in the embedding space.

**Figure supplement 8**. Optical flow estimated from FPA$_{\mathrm{uhf}}$ features and its position decoding results (**a**) Optical flow estimated from FPA$_{uhf}$ features (temporal bin size: 100 ms) showed position-dependent patterns on a circular track (Dataset 1). The vector field patterns showed trial-by-trial consistency in three representative trials. See also **video 3**. (**b**) Histograms and error CDF curves (solid line) of cross-validated decoding errors based on the optical flow of FPA$_{uhf}$ features during maze run (Dataset 1). In the CDF curve, results derived from FPA$_{uhf}$ (dotted line, same as **Fig. 1b**) are shown for comparison. (**c**) Comparison of median position decoding error in run behavior derived from different spatiotemporal features (the numbers ① ② ③ ④ refer to the same spatiotemporal feature label as in **Fig. 4**) and their combinations (Dataset 1). Error bar shows the bootstrapped SD.

**Figure supplement 9**. Comparison of discrimination accuracy in prospective position decoding (L vs. R arm) derived from clustered spikes, FPA$_{uhf}$ and dLFP$_{\theta}$ for both correct and error trials. The number of correct/incorrect trials are shown in the titles of subfigure. Results of four sessions are shown for each mouse (Dataset 6). In each session, we ran 1,000 Monte Carlo runs to compute the mean and SD statistics of 10-fold cross-validation for correct and incorrect trials. Shuffled statistics were also computed for each method and shaded area represents the SD derived from 1,000 random samples. (**a**) mouse 1. (**b**) mouse 2.

**Figure supplement 10**. Comparison of linear SVM classification accuracy (L vs. R arm) derived from clustered spikes clustered spikes, FPA$_{uhf}$ and dLFP$_{\theta}$ features for correct trials only. The number of correct trials are shown in the titles of subfigure. Results of four run sessions are shown for each mouse (Dataset 6), in both backward (*top row*) and forward (*bottom row*) directions. In each session, we ran 1,000 Monte Carlo runs to compute the mean and SD statistics of 10-fold cross-validation for correct trials. (**a**) mouse 1. (**b**) mouse 2.

**Table supplement 1.** Summary of experimental datasets under investigation. Mouse 1 consisted of five sessions, but only four sessions contained well-isolated hippocampal units.

| Dataset | Animal | # Session | Environment | # LFP channels | # sorted units |
|---|---|---|---|---|---|
| 1 | Rat 1 | 1 | circular maze (1 m diameter) | $64\times2$ channels | 92 |
| 2 | Rat 1 | 1 | linear track (1.6 m length) | $64\times2$ channels | 120 |
| 3 | Rat 2 | 1 | square T-maze | $256\times2$ channels | 223 |
| 4 | Rat 2 | 1 | linear track | $256\times2$ channels | 261 |
| 5 | Rat 3 | 1 | Open field ($120 \times 120$ $cm^2$) | $32\times2$ channels | 78 |
| 6 | Mouse 1 | 5 | circular T-maze | 64 channels | 46~59 |
| | Mouse 2 | 4 | circular T-maze | 64 channels | 12~24 |

**Table supplement 2.** Summary and comparison of clustered spike-based and LFP-based decoding methods.

| Method | Features | Likelihood | Estimator |
|---|---|---|---|
| Bayesian | clustered spikes | Poisson | Bayesian |
| OLE | clustered spikes | Gaussian | linear LS regression |
| OLE | clustered spikes | Gaussian | sparse VB-ARD linear regression |
| Likelihood | LFP-based features | Gaussian | MLE |
| Bayesian | LFP-based features | Gaussian | Bayesian |
| OLE | LFP-based features | Gaussian | linear LS regression |
| OLE | LFP-based features | Gaussian | sparse VB-ARD linear regression |

**Table supplement 3.** Result comparison between online and offline significance assessment of rat hippocampal memory replay events (Dataset 1).

| | Off-line | Online |
|---|---|---|
| # detected candidate events | 727 | 672 |
| # significant replay events | 129 | 105 |
| # significant replay events detected by both methods | 56 | |
| # significant replay events detected by off-line method only | 73 | |
| # significant replay events detected by on-line method only | 49 | |

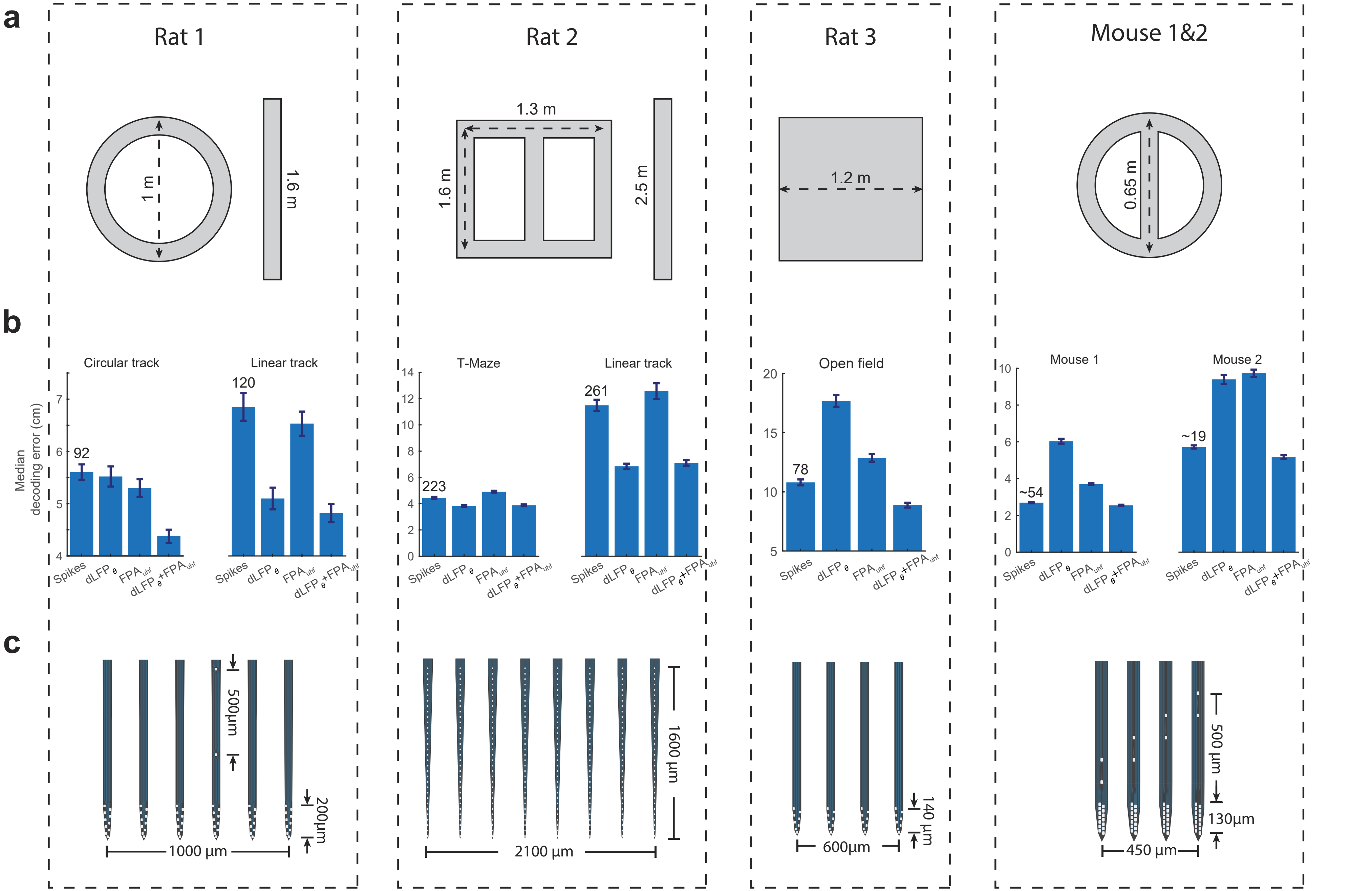


**Figure supplement 1.** Position decoding summary from different dataset with different maze and silicon probe configurations. (**a**) Schematic of four spatial environments using Datasets 1-6. (**b**) Summary of median position decoding error during maze run derived from four different features. Error bar shows the bootstrapped SD. The following default setup was used: 100 ms bin size; OLE decoding method using sparse Bayesian regression for rat 1,2 and moue 1,2; Bayesian filter decoder with a Gaussian temporal prior for rat 3. The variabilities across animals is largely part of electrode type used. $dLFP_\theta$ decoding was superior in rat 1 because large numbers of sites across all layers of the hippocampus was used but only a fraction of those were in the pyramidal layer. The number of clustered single units in each session are indicated above the spike bar. (**c**) Configuration of the recording silicon probes used in each dataset.

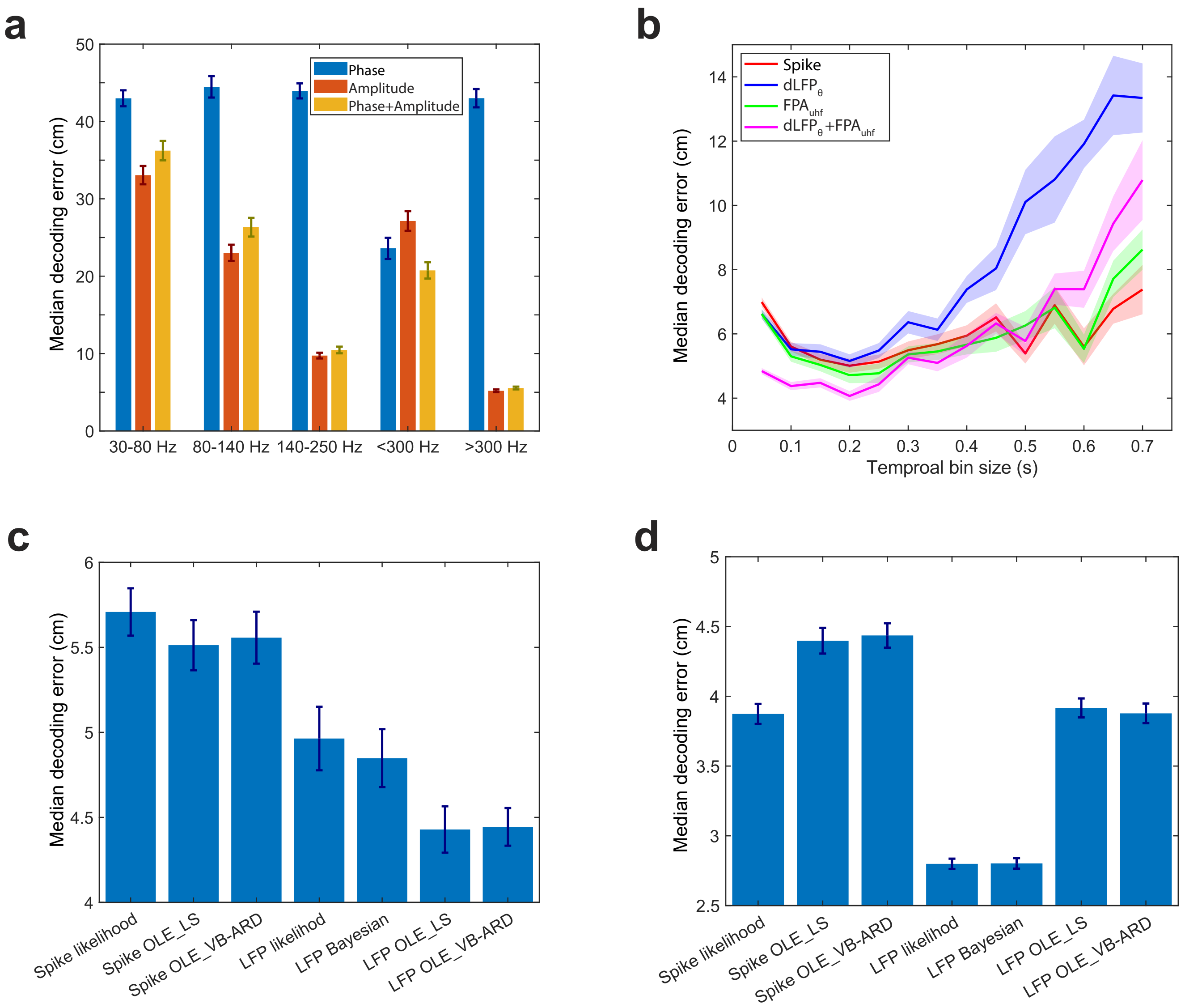


**Figure supplement 2**. Comparison of median position decoding error results derived from different band-pass filtered LFP features (phase, amplitude, phase+amplitude) at (**a**) different frequency bands, (**b**) different temporal bin size, and (**c**) different decoding methods (see **Figure 1 — table supplement 2**; Dataset 1). Shaded area or error bar shows the bootstrapped SD. (**d**) Same as (**c**), except for Dataset 3 using a different configuration of silicon probe.

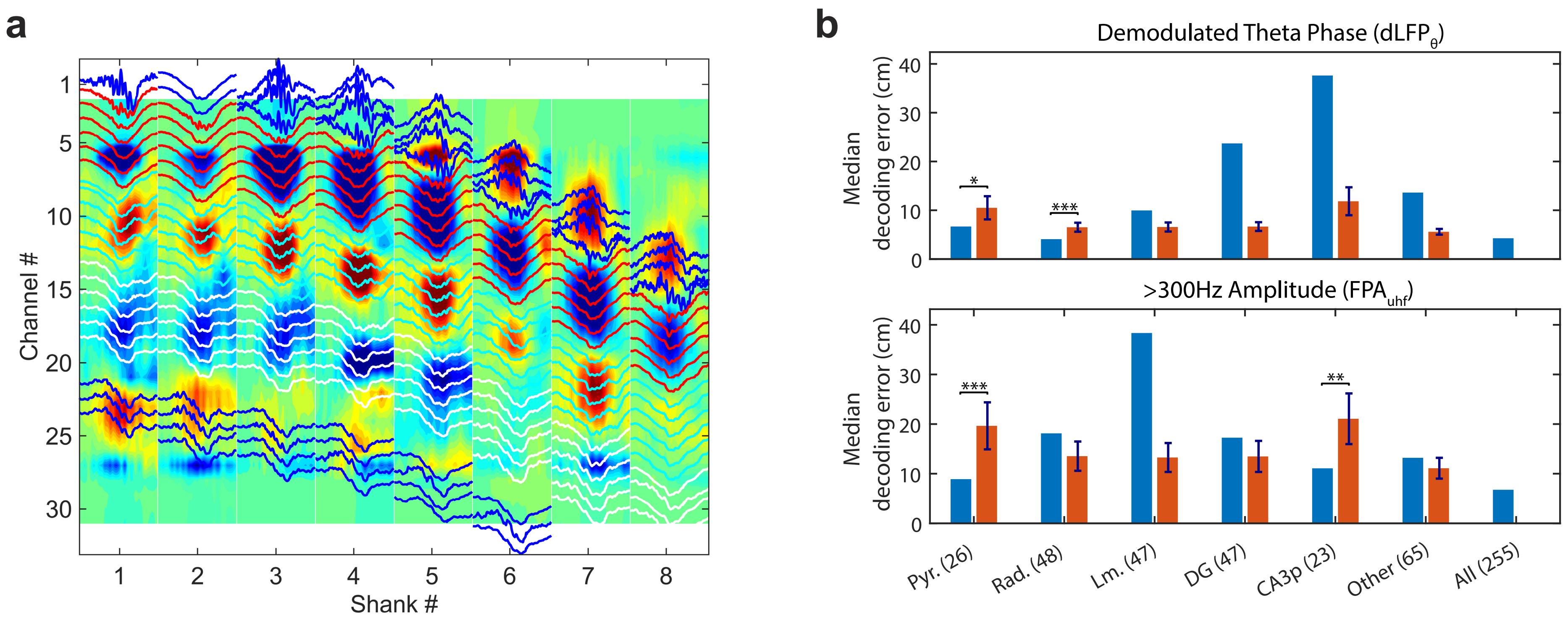


**Figure supplement 3**. Position decoding using recording sites in different hippocampal layers. (**a**) Sharp wave ripple triggered current source density (CSD) analysis revealed distinct hippocampal LFP sinks and sources in different anatomical layers (256 channels, 8 shanks, Dataset 3). (**b**) Demodulated $dLFP_{\theta}$ (*top*) and $FPA_{uhf}$ (*bottom*) median position decoding error using recording sites in different layers (*blue*) compared to median position decoding error (error bar: SD) derived from 1,000 random selections of matched number of recording sites (red). (Pyr: stratum pyramidale; Rad: stratum radiatum; Lm: stratum lacunosum moleculare; DG: dental gyrus; CA3p: CA3 pyramidal layer). The number inside parentheses indicates the channel count within that layer/region. CA1 layers with changing layer-by-layer theta phase shift gave best position decoding by $dLFP_{\theta}$ decoding strategy, while $FPA_{uhf}$ was most effective in CA1 and CA3 pyramidal layer.

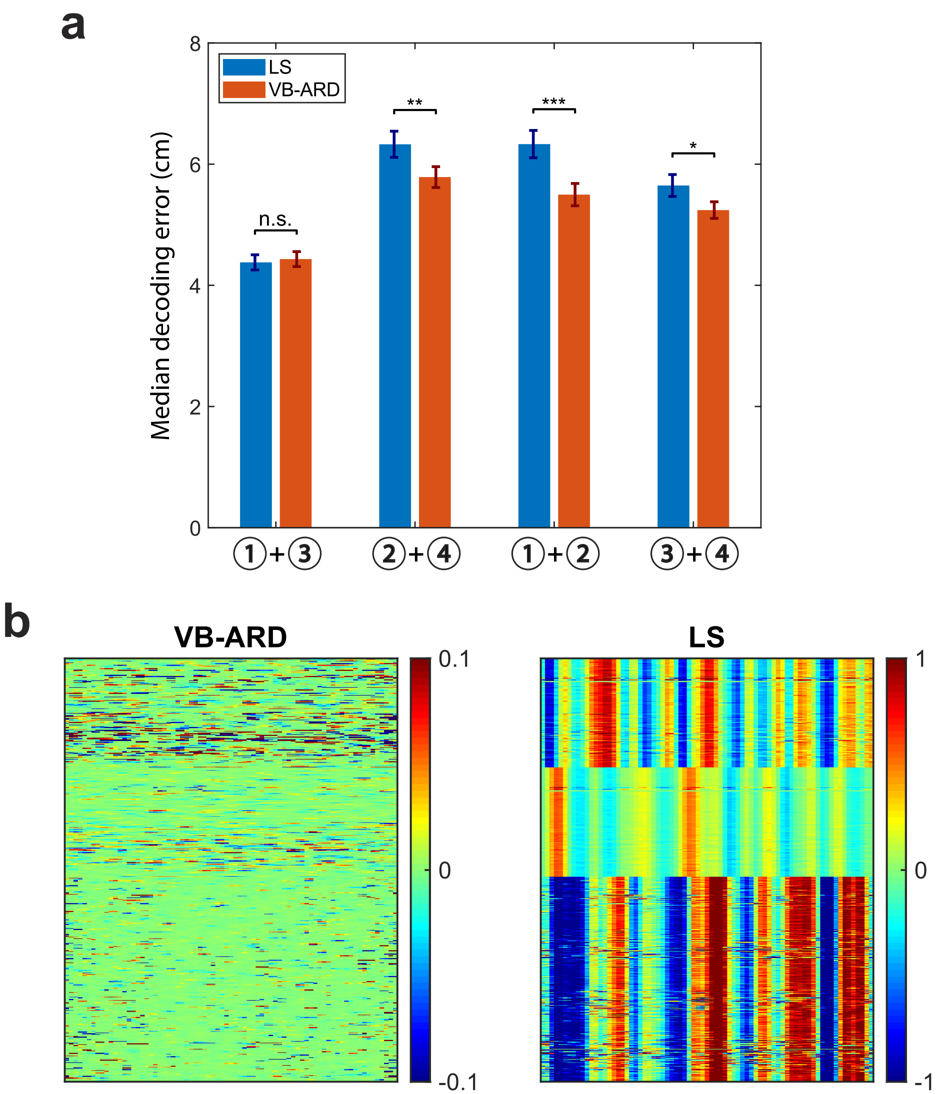


**Figure supplement 4**. (**a**) Comparison of median decoding error results (Dataset 1) between methods using standard least-squared (LS, blue) and sparse VB-ARD (red) regression while testing different combinations of LFP-based features (①: dLFP$_{\theta}$ amplitude and phase; ②: optical flow at 4-12 Hz band; ③: >300 Hz amplitude (FPA$_{uhf}$); ④: optical flow at >300 Hz band). *, P<0.05; **, P<0.01; *** P<0.001, rank-sum test. Error bar shows the bootstrapped SD. (**b**) Comparison of the mapping *V* between the VB-ARD (*left*) and LS (*right*) estimators used in the OLE method ("①+②" LFP features combination). The weight coefficients learned from the VB-ARD estimator were sparse.

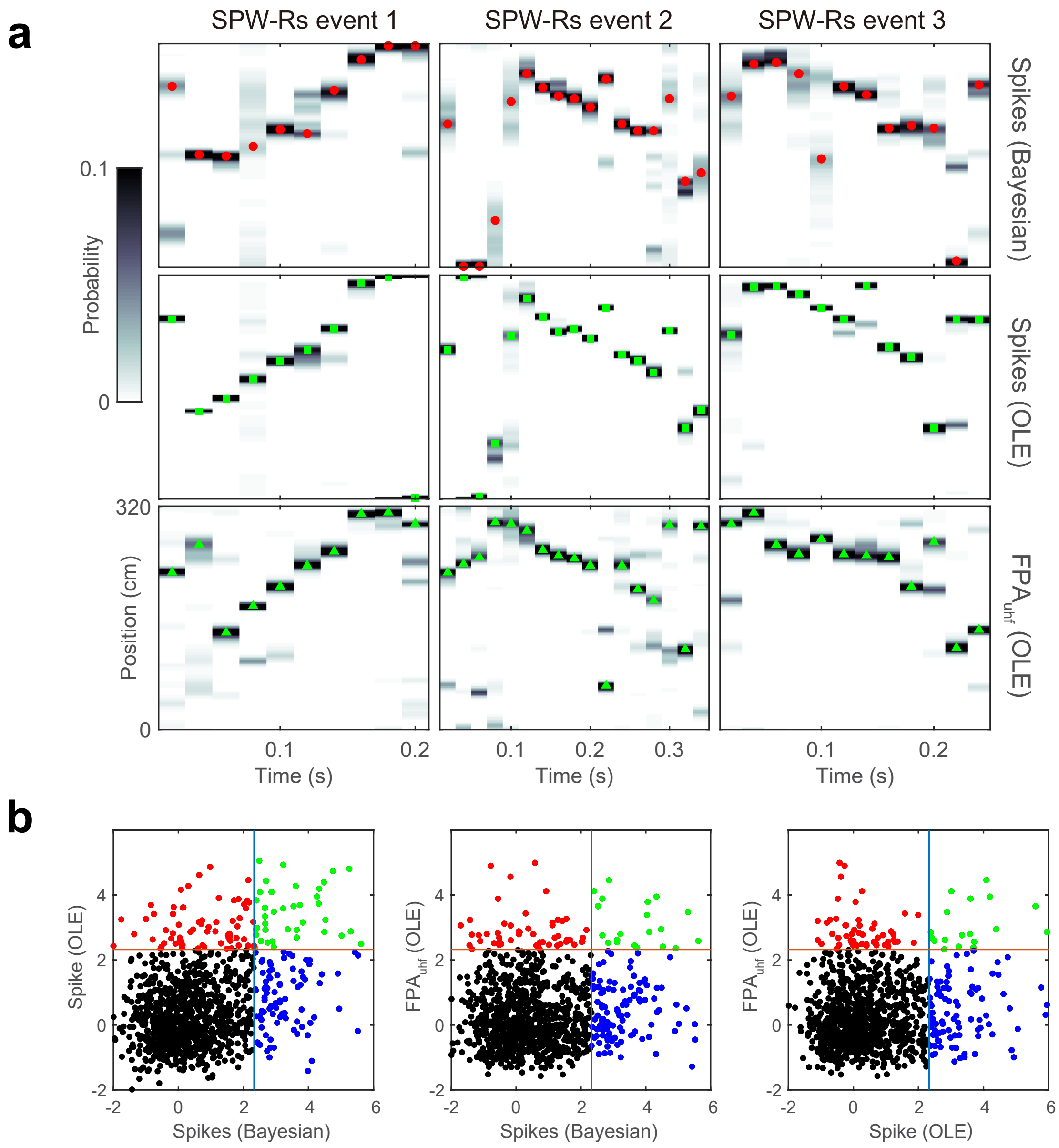


**Figure supplement 5**. Examples of decoded ripple-replay contents derived from 2nd dataset. (**a**) Examples of decoded ripple-replay contents derived from clustered spikes and $FPA_{uhf}$ features in post-run sleep (Datasets 2). (**b**) Comparison of detection statistics of hippocampal memory replay candidates in awake and NREM ripples (n=1,038 candidate events) derived from clustered spikes and $FPA_{uhf}$ features.

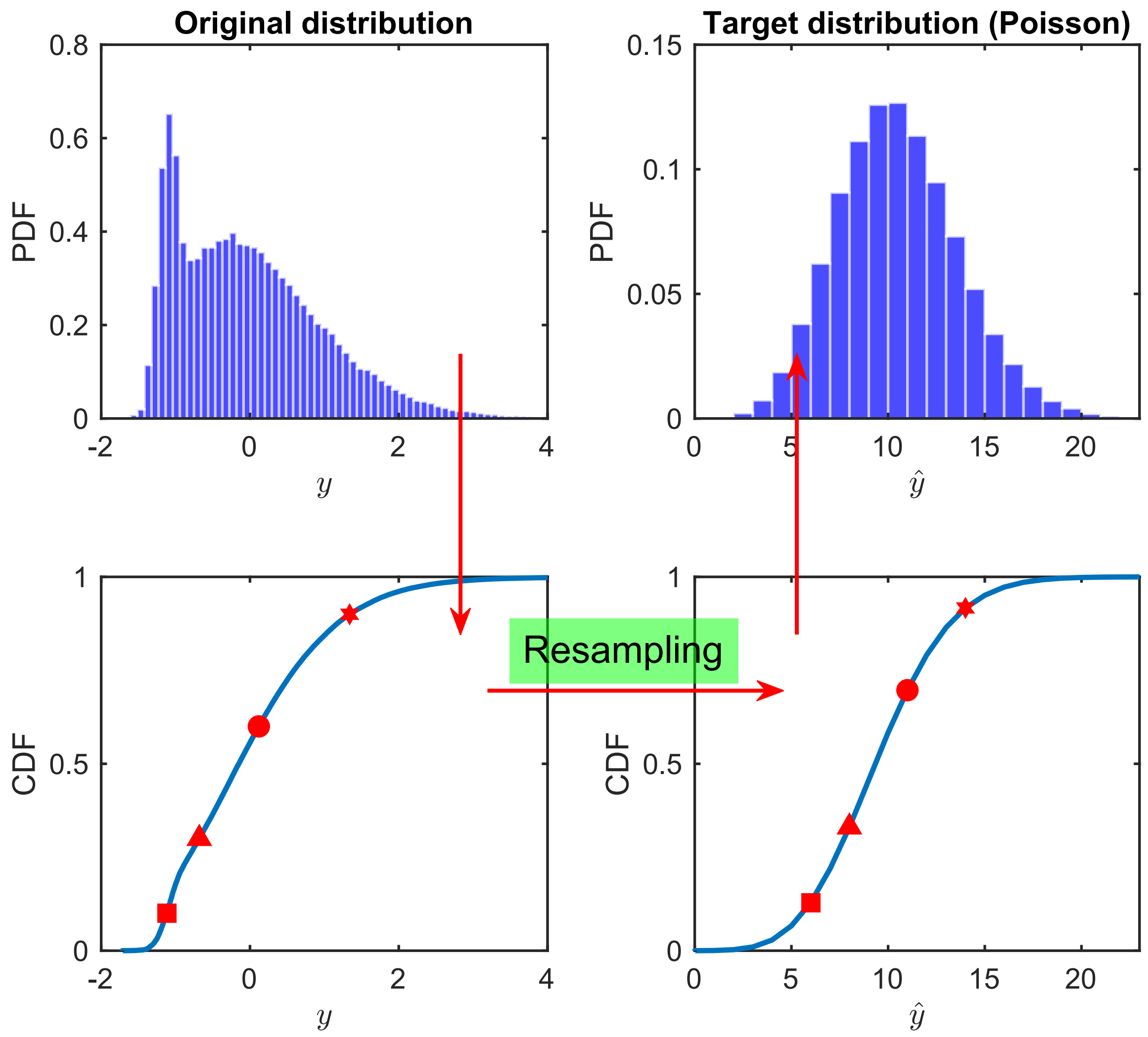


**Figure supplement 6**. Schematic procedure of resampling to rescale the Z-scored real-valued observations $y$ (from an empirical distribution) to nonnegative count observations $\hat{y}$ (from a target Poisson distribution with mean statistic of 10). The red arrows indicate the conceptual procedure. Note that the order statistics remained unchanged between two cumulative distributed function (CDF) curves.

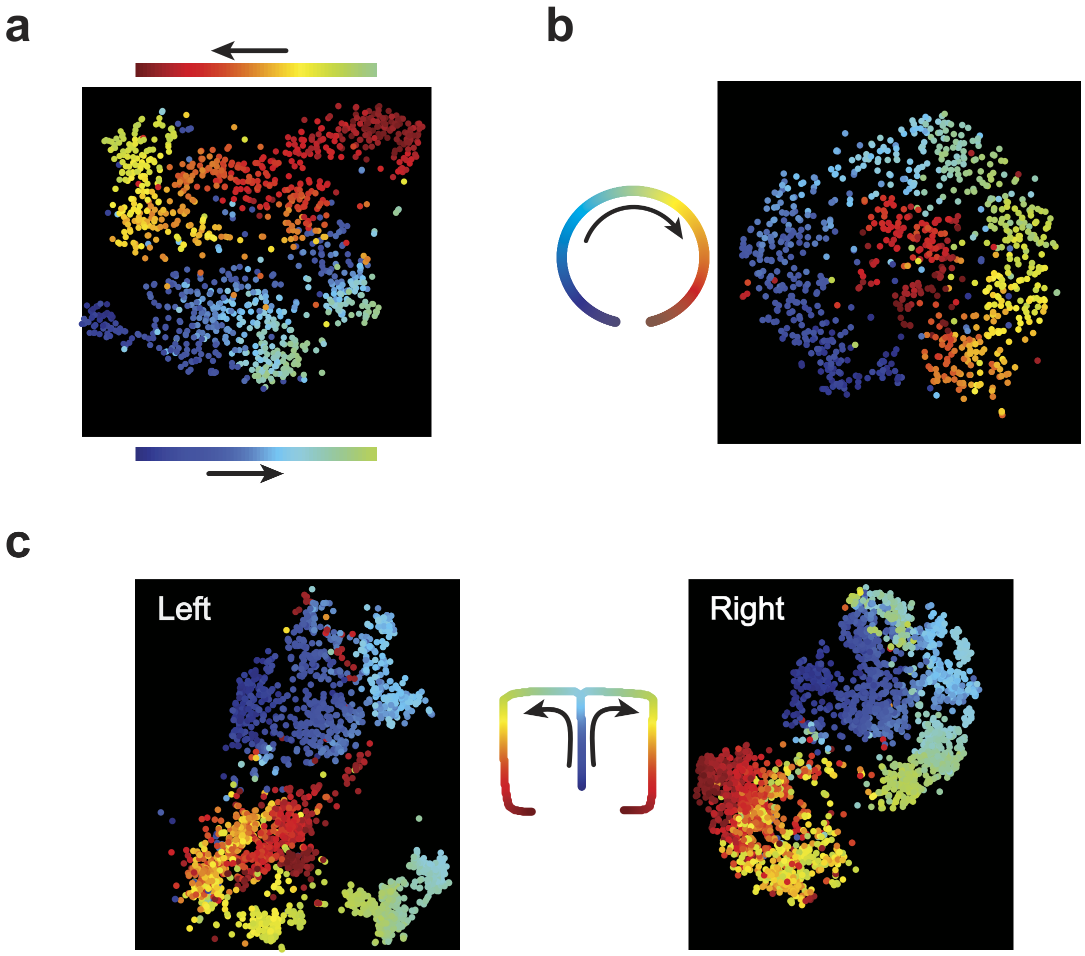


**Figure supplement 7.** Dimensionality reduction of multi-site recorded $FPA_{uhf}$ features during track running, produced by t-distributed stochastic neighbor embedding (t-SNE) algorithm. Each dot was color-coded in relation to the animal's position on the track. Two multi-site high-dimensional $FPA_{uhf}$ features were similar if they were close in the two-dimensional embedding space. (**a**) For the circular track (Dataset 1), we recovered a continuous trajectory pattern in the embedding space. (**b**) For the linear track (Dataset 2), we recovered direction-specific trajectory pattern in the embedding space. (**c**) For the T-maze (Dataset 3), we recovered distinct clustered patterns, corresponding to L and R turns, in the embedding space.

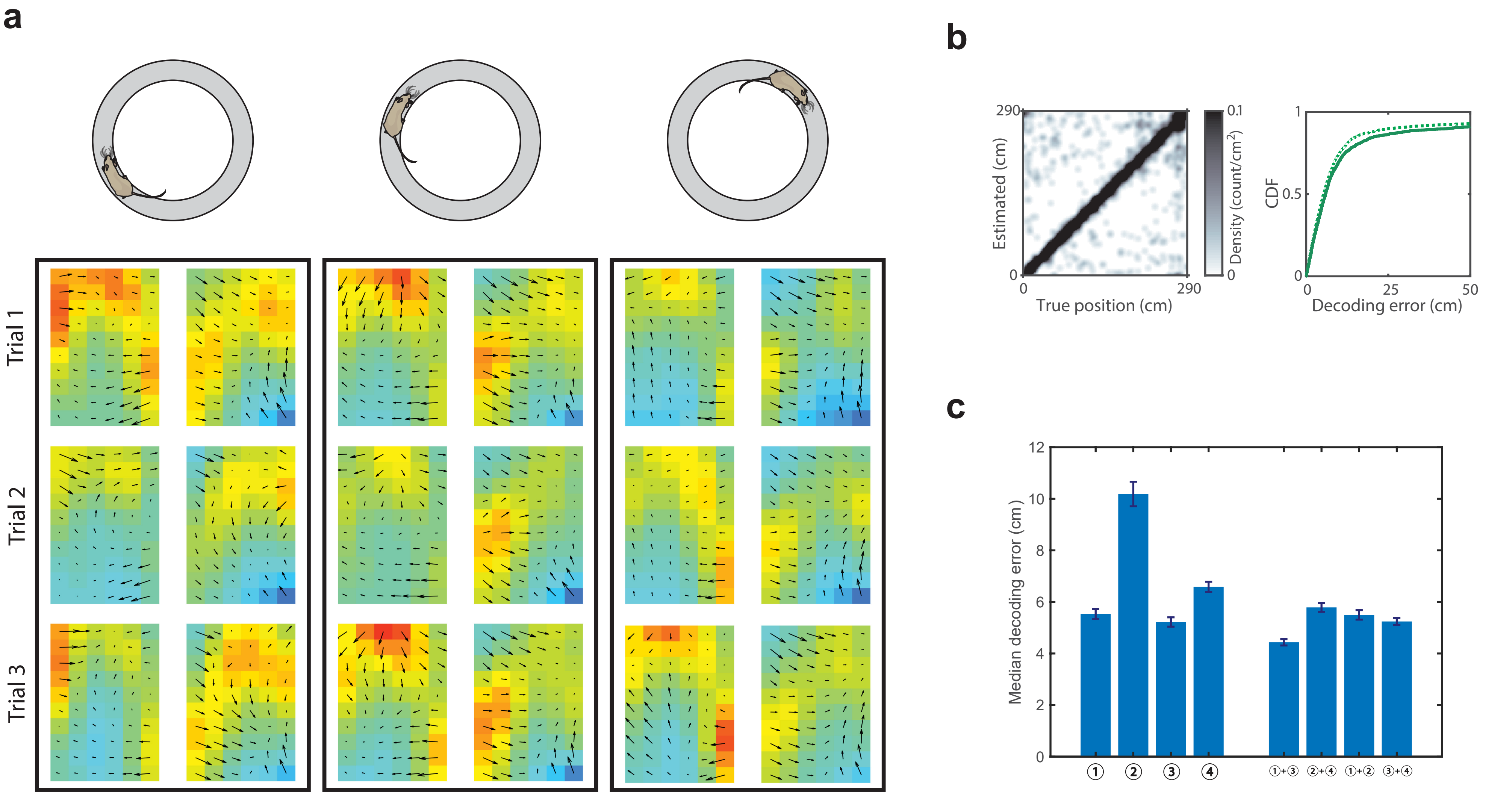


**Figure supplement 8.** Optical flow estimated from $FPA_{uhf}$ features and its position decoding results (**a**) Optical flow estimated from $FPA_{uhf}$ features (temporal bin size: 100 ms) showed position-dependent patterns on a circular track (Dataset 1). The vector field patterns showed trial-by-trial consistency in three representative trials. See also **video 3**. (**b**) Histograms and error CDF curves (solid line) of cross-validated decoding errors based on the optical flow of $FPA_{uhf}$ features during maze run (Dataset 1). In the CDF curve, results derived from $FPA_{uhf}$ (dotted line, same as Fig. 1b) are shown for comparison. (**c**) Comparison of median position decoding error in run behavior derived from different spatiotemporal features (the numbers ① ② ③ ④ refer to the same spatiotemporal feature label as in Fig. 4) and their combinations (Dataset 1). Error bar shows the bootstrapped SD.

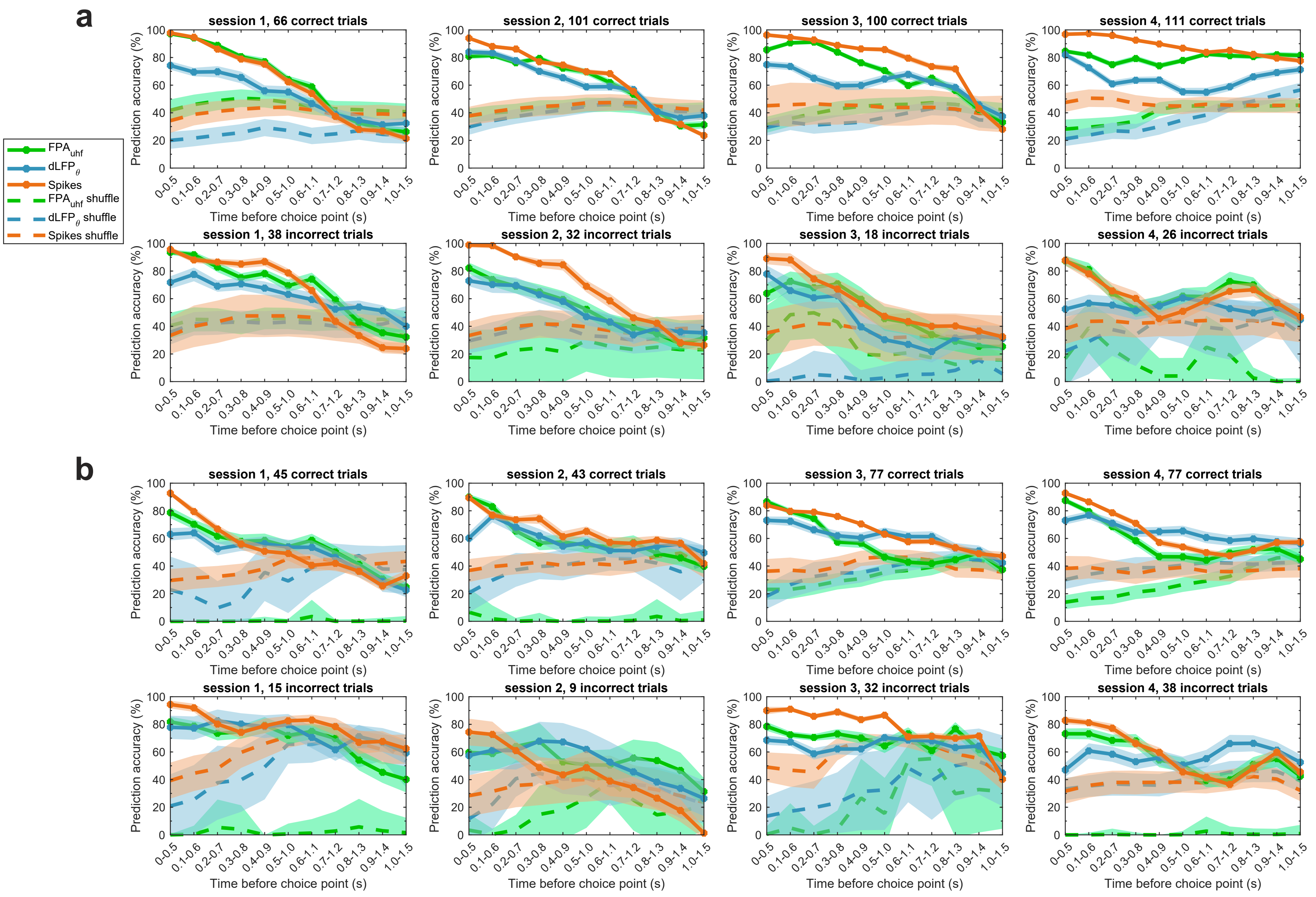


**Figure supplement 9**. Comparison of discrimination accuracy in prospective position decoding (L vs. R arm) derived from clustered spikes, FPA$_{uhf}$ and dLFP$_{\theta}$ for both correct and error trials. The number of correct/incorrect trials are shown in the titles of subfigure. Results of four sessions are shown for each mouse (Dataset 6). In each session, we ran 1,000 Monte Carlo runs to compute the mean and SD statistics of 10-fold cross-validation for correct and incorrect trials. Shuffled statistics were also computed for each method and shaded area represents the SD derived from 1,000 random samples. (**a**) mouse 1. (**b**) mouse 2.

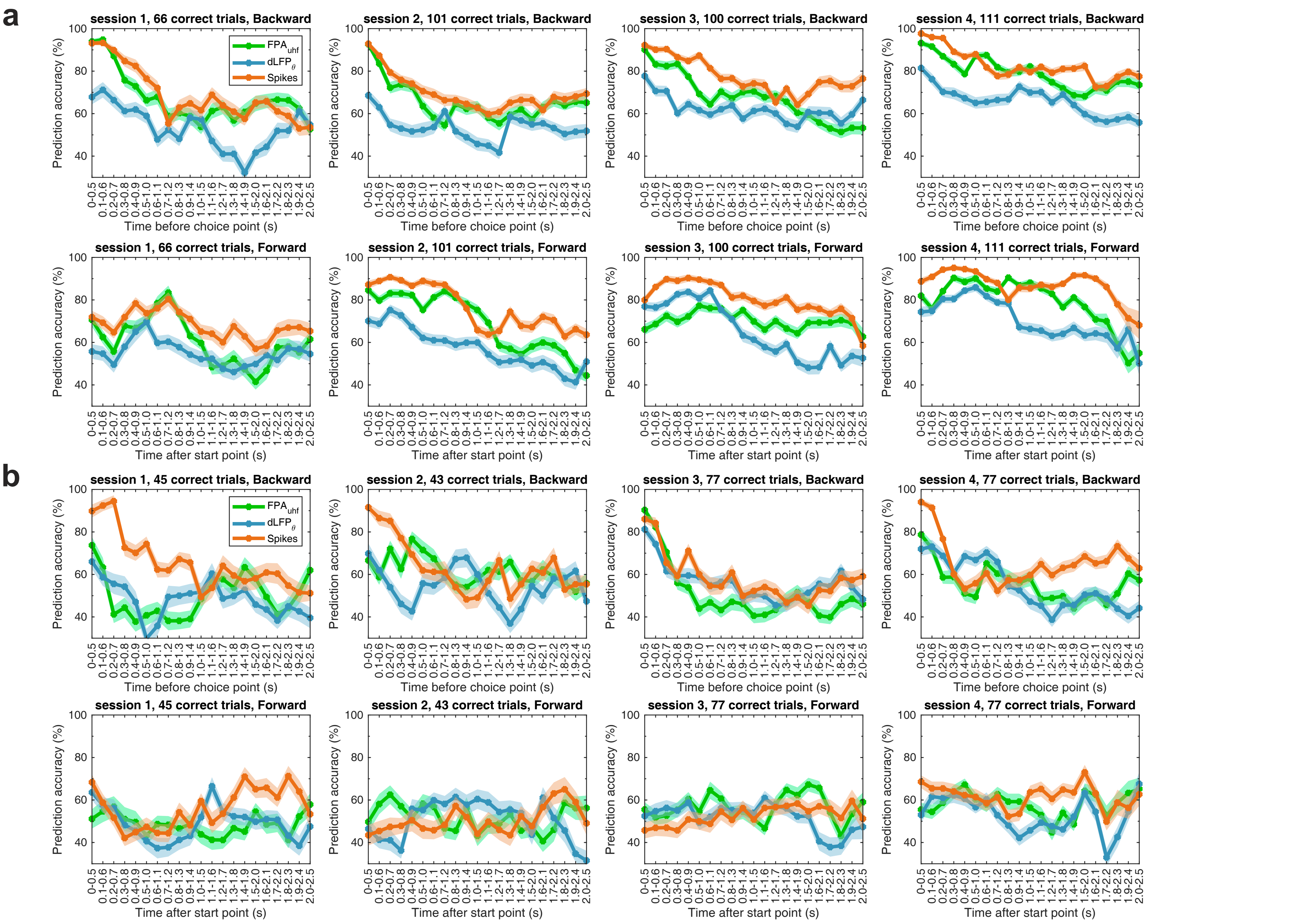


**Figure supplement 10.** Comparison of linear SVM classification accuracy (L vs. R arm) derived from clustered spikes clustered spikes, FPA$_{uhf}$ and dLFP$_{\theta}$ features for correct trials only. The number of corrrect trials are shown in the titles of subfigure. Results of four run sessions are shown for each mouse (Dataset 6), in both backward (top row) and forward (bottom row) directions. In each session, we ran 1,000 Monte Carlo runs to compute the mean and SD statistics of 10-fold cross-validation for correct trials. (**a**) mouse 1. (**b**) mouse 2.
